# Interpretable scRNA-seq Analysis with Intelligent Gene Selection

**DOI:** 10.1101/2024.09.01.610665

**Authors:** Tianhao Ni, Xinyu Zhang, Kaixiu Jin, Guanxiong Pei, Nan Xue, Guanao Yan, Taihao Li, Bingjie Li

## Abstract

Single-cell RNA sequencing (scRNA-seq) data analysis faces multiple challenges, including high dimensionality, significant noise, and data loss. To effectively address these issues, we introduce AIGS, a robust and transparent single-cell analysis framework. AIGS utilizes an intelligent gene selection method that systematically identifies the most informative genes for clustering based on the normalized mutual information between pre-learned pseudo-labels and quantified genes. Additionally, AIGS incorporates a scale-invariant distance metric to assess cell-to-cell similarity, enhancing connections between homogenous cells and ensuring more accurate and robust results. Through comprehensive comparisons with state-of-the-art techniques, AIGS demonstrates superior performance in both clustering accuracy and multi-resolution visualization quality. Our in-depth analysis of clustering and visualization results further reveals that AIGS can uncover complex, stage-specific gene expression patterns during the same developmental cell stage.

## 1 Introduction

The recent rise of single-cell RNA sequencing (scRNA-seq) has enabled the detection of cell types at the molecular level. Cell annotation, also known as cell clustering, is a crucial step in scRNA-seq analysis. It enables the identification of cell populations [1], determination of cell population topology [2], and characterization of cellular heterogeneity in complex diseases [3]. The main challenges of clustering come from two aspects. The first challenge arises from the extremely high dimensionality of the RNA-seq data, which presents obstacles in accurately characterizing the spatial distribution of data using conventional distances such as Euclidean and cosine distance. The second challenge arises from the technical limitations of current sequencing methods, leading to missing gene expression readings (dropouts) and outliers [4]. While classical single-cell analysis frameworks like Seurat [5] and SIMLR [6] are widely used, they may fall short of achieving high clustering accuracy. Deep network clustering approaches like scDHA [7] have shown significant improvements in performance due to their extreme representational power. Nonetheless, deep networks are commonly perceived as black-box models, and the mechanism of extracting clustering-relevant information from high-dimensional data remains unclear, which hinders the further development of deep networks.

To overcome these challenges, we present AIGS (single cell Analyzer with Intelligent Gene Selection), an interpretable framework designed for accurate and efficient scRNA-seq analysis. The AIGS pipeline comprises modules for gene selection, dimensionality reduction, clustering, visualization, and marker gene identification. AIGS distinguishes itself from other frameworks by utilizing an intelligent gene selection algorithm that targets genes which indicate cell types, a minority of all genes that provide the most informative data on cell types. This gene selector systematically identifies class-indicating genes based on the normalized mutual information (NMI) between the learned pseudo-labels and quantified genes, effectively reducing data dimensionality and mitigating the negative impact of dropouts. Furthermore, AIGS incorporates a novel scale-invariant distance metric that highlights the similarity among homogeneous cells while differentiating heterogeneous cells. Notably, AIGS emphasizes functional clustering, wherein cells with similar functions or states are grouped together, yielding deeper insights into the underlying biological processes. This combination of features establishes AIGS as a promising framework for the analysis of scRNA-seq data. And furthermore, AIGS’s outstanding performance does not rely on unpredictable outcomes from deep learning frameworks, making it applicable to other analytical tasks, including spatial transcriptomics and multimodal single-cell genomics.

## 2 Results

### 2.1 A brief overview of AIGS

AIGS is a versatile cell data analysis framework that accommodates various data types and relies on general machine learning algorithms, ensuring robust interpretability (**Fig. 1a**). One of its pioneering features is an intelligent gene selection module, which plays a pivotal role in achieving precise cell analysis. This module strategically chooses a minimal set of genes while encompassing nearly all marker genes. This strategy significantly enhances analysis efficiency and conserves computational resources, effectively addressing the challenge posed by ultra-high dimensionality in cell data.

**Figure 1:**
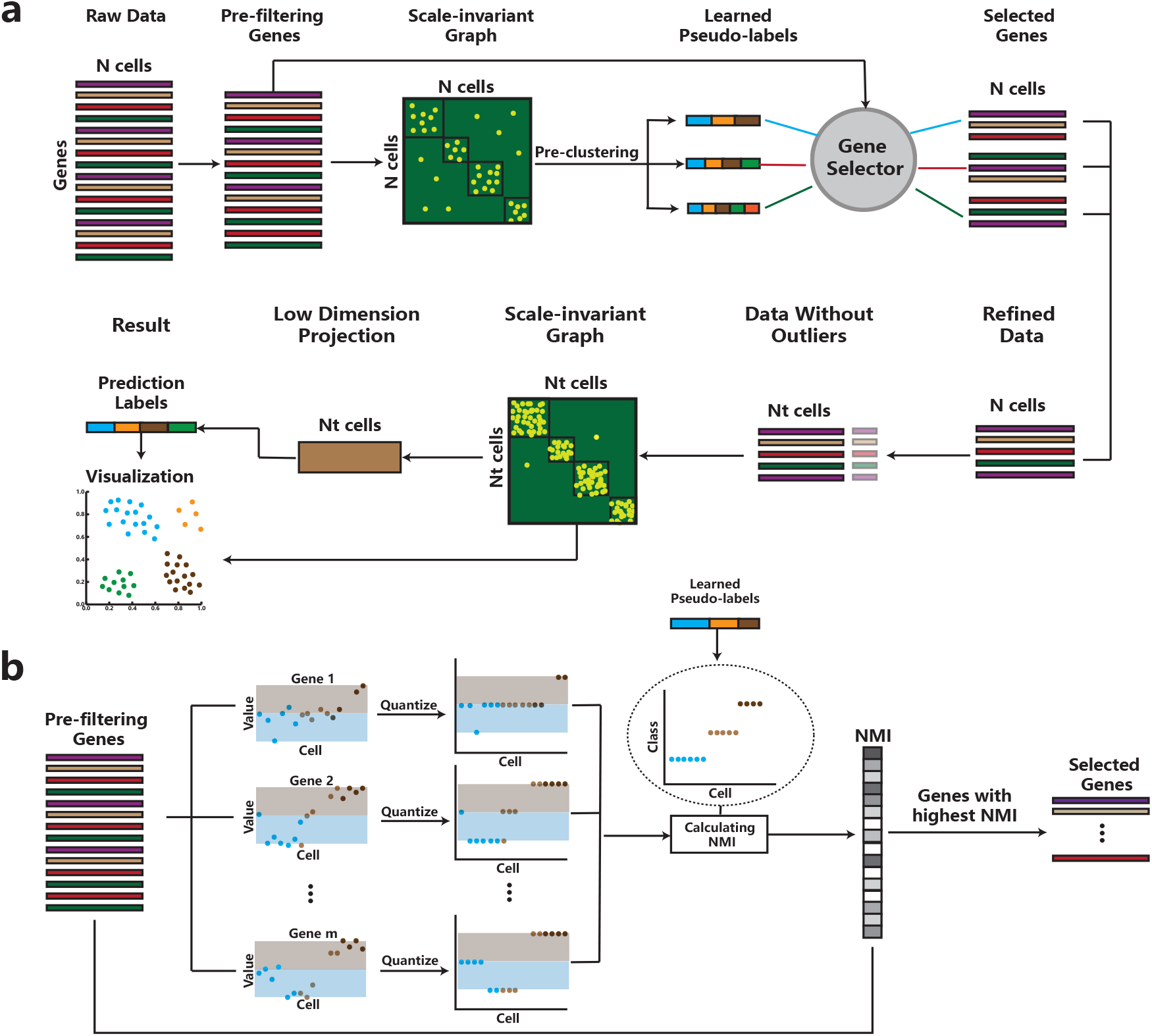
The AIGS framework for analyzing scRNA-seq data. (a) An overview of the AIGS pipeline. Initially, AIGS undertakes preprocessing of the gene expression matrix data, subsequently constructing a similarity graph utilizing a scale-invariant metric for quantifying cell-to-cell relationships. Upon this graph, AIGS selects class-indicating genes, detects and removes outlier cells, estimates the number of cell classes, and performs low-dimensional projection and clustering. The framework also provides cell visualization based on the similarity graph and clustering results, employing varying color schemes to signify disparate cell classifications. (b) An illustration of the gene selector of the AIGS pipeline. The gene selector learns the pseudo-label and calculates the normalized mutual information (NMI) between the pseudo-label and the quantified genes. It shortlists a minimal proportion of genes (default as 100) with the highest NMI and takes the union of selected genes from multiple pseudo-labels. For instance, the quantified results of gene 1 and gene 2 exhibit consistency with a three-class pseudo-label and are therefore selected, while that of gene *m* have consistent values across most cells and are consequently not chosen by AIGS.

For instance, in the context of single-cell RNA sequencing data, initiates the process by preprocessing the cell data, excluding genes with abnormal expression levels attributable to experimental errors. Subsequently, it constructs a scale-invariance similarity graph, where nodes represent cells, and edge weights signify the likelihood of cell homogeneity. The framework employs spectral embedding [8] to project this graph into a low-dimensional space and applies K-means clustering [9] to derive pseudo-labels for the low-dimensional data. Using these pseudo-labels, AIGS further selects representative genes for each cluster.

Specifically, the gene selection process is guided by quantization and normalized mutual information metrics (NMI, **Supplementary Note 4**). AIGS quantifies each gene and computes NMI with pseudo-labels. The top 100 genes, exhibiting the highest similarity with pseudo-labels, are chosen. Genes that display distinctive expression within one cluster, setting them apart from others, are signed as selected genes (like gene 1 and gene 2 in **Fig. 1b**). Conversely, genes with similar expression across all clusters are excluded (like gene m in **Fig. 1b**). This gene selection methodology effectively identifies marker genes for specific clusters or cell subtypes, thereby standardizing similarity criteria among cells and enabling the clear delineation of heterogeneous cells while closely grouping homogeneous ones.

Following gene selection, AIGS measures the similarity between each cell and its neighboring cells within its local neighborhood. It eliminates the most distant cells from their neighborhoods, mitigating technical noise stemming from amplification biases, batch effects, and sequencing errors.

AIGS proceeds by reconstructing a new scale-invariance graph for the processed data, applying spectral embedding for dimensionality reduction, and utilizing K-means clustering. The optimal number of clusters is determined based on the consistency between clustering results and the adjacency matrix of the graph. Subsequently, a graph-based visualization algorithm generates visualizations at multiple resolutions, and marker genes for each cell type in the clustering and visualization results are identified using analysis of variance.

In summary, AIGS offers a comprehensive and efficient analysis pipeline encompassing gene selection, outlier cell removing, and a range of downstream analyses, including clustering, visualization, subtyping, and marker gene selection. Notably, these steps rely solely on computation of cell-to-cell distances, making AIGS applicable to various single-cell tasks across different data types.

### 2.2 AIGS enables the detection of cell-type-related genes

The gene selection process in single-cell RNA sequencing (scRNA-seq) analysis is crucial, as it significantly influences the quality and accuracy of subsequent analyses. Unlike some other scRNA-seq analysis methods like Seurat[5], which retain a large number of genes for analysis, AIGS takes a unique approach. AIGS constructs pseudo-labels and selectively retains a smaller yet more informative set of genes, typically in the range of 100 to 300 genes (**Fig. 2a** and **Supplementary Fig. S1**).

**Figure 2:**
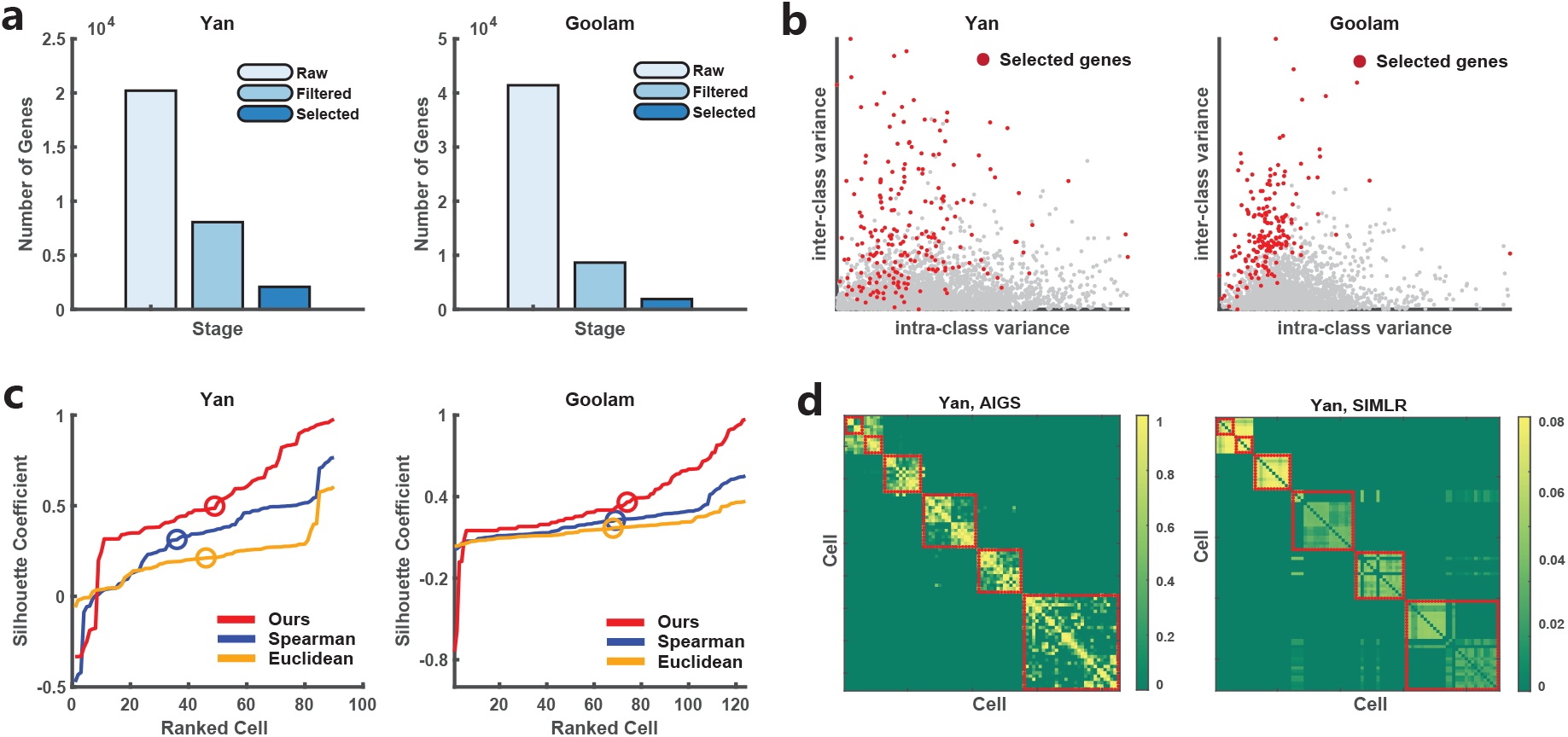
Advantages of gene selection and scale-invariant distance in the AIGS framework demonstrated on the Yan [12] and Goolam [13] datasets.(a) Comparison bar chart of retained gene counts in different stages of AIGS. From left to right, Raw data, after gene filtering, after gene selection. (b) Comparison of selected and eliminated genes’ intra-class and inter-class variances in two datasets. The red points represent genes selected by AIGS’s gene selection module, while the gray points represent genes that were not selected. (c) Comparison of silhouette coefficient distributions for Spearman distance, euclidean distance, and scale-invariant Distance, with circles representing the average silhouette coefficients for each distance. (d) Comparison of the similarity matrices constructed by AIGS and SIMLR on the Yan dataset. The cells within the red boxes represent the intra-class cell similarities that match the golden label, while the cells outside the boxes represent inter-class cell similarities.

A key feature of AIGS’s gene selection strategy is its focus on class-indicative properties. This means that the genes chosen by AIGS tend to exhibit consistent expression levels within the same cell type while showing distinct expression patterns among different cell types. This selectivity ensures that downstream analyses are driven by genes that are highly informative for characterizing cell types and their differences.

To validate the effectiveness of AIGS’s gene selection process, experiments were conducted on various datasets, including the Yan [12] and Goolam [13] datasets, as well as six others [14, 15, 16, 17, 18, 19] (**Supplementary Fig. S19** and **Supplenmentary Table 1**). The assessment primarily focused on measuring the intra-class variance (variance within the same cell type) and inter-class variance (variance between different cell types) of gene expression levels (**Fig. 2b** and **Supplementary Fig. S2**). The effect described can also be intuitively illustrated through t-SNE visualizations, both before and after gene selection (**Supplementary Fig. S3** and **Fig. S4**). In the visualization results, the gene selection module not only brings together cells of the same class but also notably improves the distinctiveness between cells of different classes. AIGS selected genes consistently demonstrated excellent intra-class consistency and significant inter-class differences across all eight datasets.

Overall, AIGS’s gene selection strategy, which retains a compact set of class-indicative genes, ensures that the selected genes not only represent cell types accurately but also distinguish between them effectively. This approach ultimately enhances the precision and reliability of downstream analyses in scRNA-seq data.

### 2.3 AIGS effectively improves the spatial distribution of scRNA-seq data

AIGS has achieved outstanding performance, with one of its core strengths being its innovative definition of scale-invariant distance. This distance measure defines the proximity between two cells based on the ranking of Spearman distances from one cell to another. In simpler terms, if two cells are each other’s closest neighbors, their distance is defined as (1).

This metric’s uniqueness lies in its sole consideration of the proximity between cells, disregarding the numerical values of distances. As a result, it remains robust even in high-dimensional and spatially complex datasets, accurately reflecting the relative proximity of cells in such spaces. In practical cell classification results, cells belonging to the same category are closer to each other under this metric, while cells from different categories are farther apart.

To illustrate the effectiveness of this distance, we applied it to the Yan and Goolam datasets. We calculated silhouette scores for each cell in the dataset using three different distance metrics (scale-invariant distance, Spearman distance, and Euclidean distance) and sorted them in ascending order (**Fig. 2c**). The scale-invariant metric outperformed the others by more accurately capturing the intricate high-dimensional distribution of cells, aligning better with the true labels. Statistical analysis, including Wilcoxon signed-rank tests, demonstrated the significant superiority of the scale-invariant metric over the others, with p-values of 2.2 × 10^−6^ and 7.8 × 10^−7^, respectively.

Furthermore, AIGS demonstrates proficiency in acquiring more refined and sparser similarity matrices through scale-invariant distance. As a result, the similarity matrix exhibits increased intra-class similarity and decreased inter-class similarity. This outcome stems from the sparse property of the scale-invariant distance calculation, where only distances between each cell and its 10 nearest neighbors are considered, while the remaining distances are set to infinity. When comparing with another graph-based learning algorithm, such as SIMLR, AIGS has the capability to learn similarities that align more accurately with genuine cell labels (**Fig. 2d** and **Supplementary Fig. S5**).

### 2.4 AIGS significantly improves the accuracy and stability of cell annotation

Upon assessing five clustering methods [5, 6, 7, 10, 11] on eight publicly available datasets [12, 13, 14, 15, 16, 17, 18, 19] (**Supplementary Table 1** and **Note 6**), we discovered that AIGS consistently exhibits superior clustering performance, exhibiting both high accuracy and reliability. We primarily used the Adjusted Rand Index (ARI) as the evaluation criterion and also considered four other widely used clustering criteria (**Supplementary Note 4**). AIGS achieved the highest rank according to ARI, with an average score of 92%, significantly outperforming the second-best method, scDHA, which attained an 83% average score (**Fig. 3a**). Furthermore, AIGS outperformed scDHA by at least 6% on average across other evaluation criteria and achieved the top ranking on most datasets (**Supplementary Table 2**). The one-sided Wilcoxon signed rank test shows that the clustering metric values of AIGS are significantly higher than other methods, with a p-value of 7.2 × 10^−7^. In other accuracy metrics, including ACC, NMI, Jaccard coefficient, and Fmeasure (**Supplementary Fig. S19**), it is equally evident that AIGS demonstrates outstanding performance in clustering tasks (**Supplementary Fig. S6** and **Table 2**).

**Figure 3:**
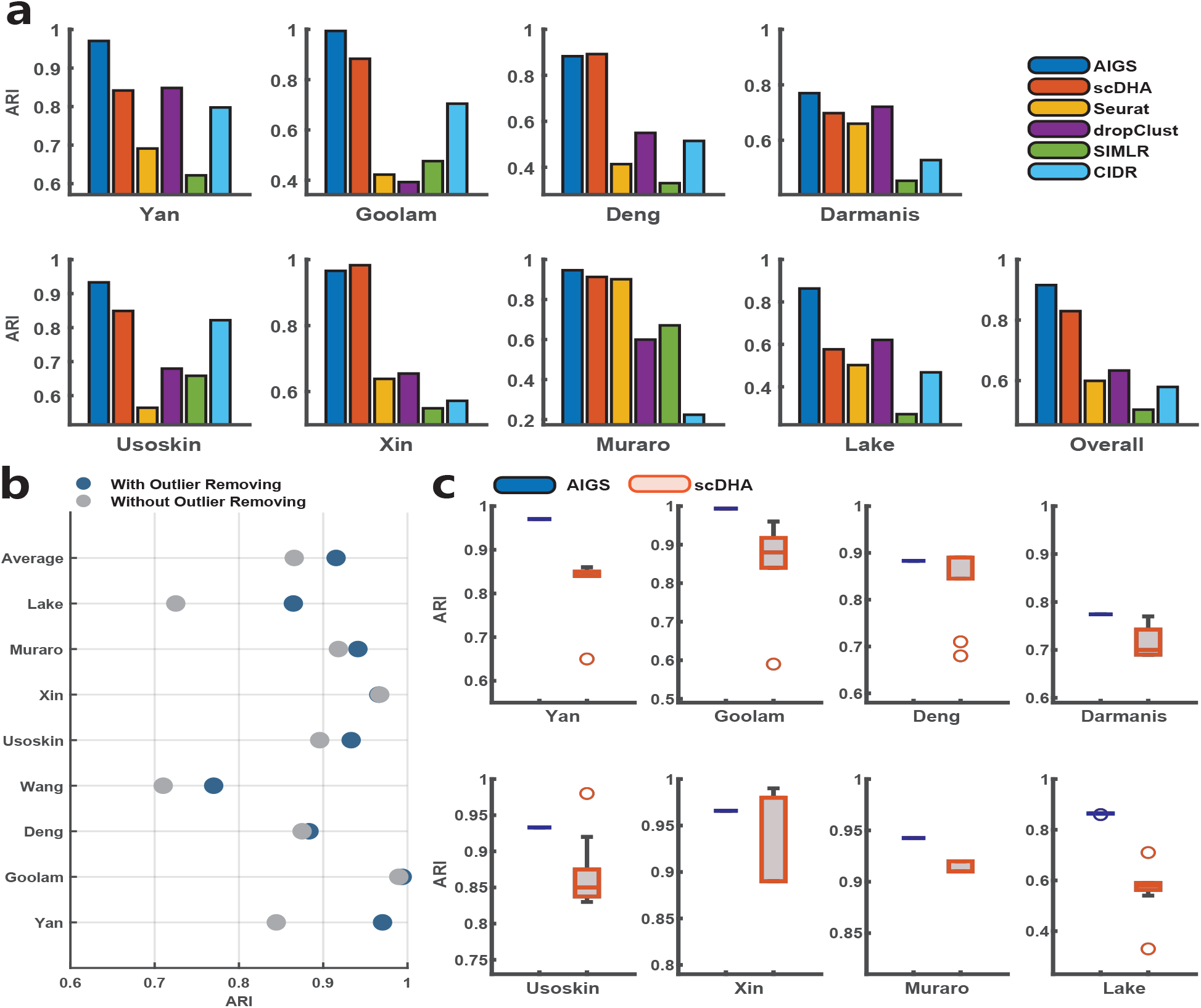
Performance comparison between our proposed AIGS and existing methods across eight published datasets [12, 13, 14, 15, 16, 17, 18, 19]. (a) The comparison bar chart of clustering accuracy (ARI) for the six methods, from left to right: AIGS, scDHA, Seurat, dropclust, SIMLR, and CIDR. (b) The impact of outlier removal on the enhancement of clustering accuracy (ARI). (c) Stability assessment of clustering performance between AIGS and scDHA, executed ten times on the specified datasets.

Furthermore, AIGS’s intelligent gene selection module and outlier cell removal module significantly enhance clustering accuracy and stability. The experiments verified that under very limited runtime conditions, the gene selection process averaged an improvement of 26.4% in clustering accuracy across eight datasets (**Supplementary Fig. S7**). One key factor is that the selected genes facilitated the normalization of similarities between samples, enabling heterogeneous cells to be distinctly partitioned and homogeneous cells to be more closely grouped. The exclusion of outliers averted the potential distortion of clustering by aberrant cells, which could form separate classes or improperly link two unrelated classes. Following outlier exclusion, the average clustering accuracy saw a further increase by 4.97% (**Fig. 3b**). The one-sided Wilcoxon signed rank test corroborates the significant improvement in clustering metric values post outlier exclusion, with a p-value of 1.8 ×10^−8^. Additionally, AIGS exhibited superior stability compared to scDHA, the second-best method (**Fig. 3c**). This is also attributed to the gene selection and abnormal cell elimination modules, which enhance the intra-class cohesion and inter-class dispersion of spectral projections, thus endowing robustness to the K-means clustering algorithm applied to the spectral projection results, significantly mitigating AIGS’s randomness. Overall, our findings demonstrate the impressive performance of AIGS in accurate and reliable scRNA-seq clustering.

### 2.5 AIGS estimates the number of cell types in an optimal manner

The AIGS framework offers a dependable approach for determining the number of cell types. Through examining diverse potential clustering patterns and gauging their aptitude to represent the inherent data structure based on cell similarity (**Methods**), AIGS always chooses the number of cell types that correspond to the highest or near-highest ARI. As demonstrated in **Fig. 4a**, we have compared the clustering performance (ARI) under different numbers of clusters, with the red line designating the number of clusters as estimated by AIGS, generally leading or closely following the best. This capability is particularly beneficial when the true number of cell types within a dataset is unknown or challenging to discern.

**Figure 4:**
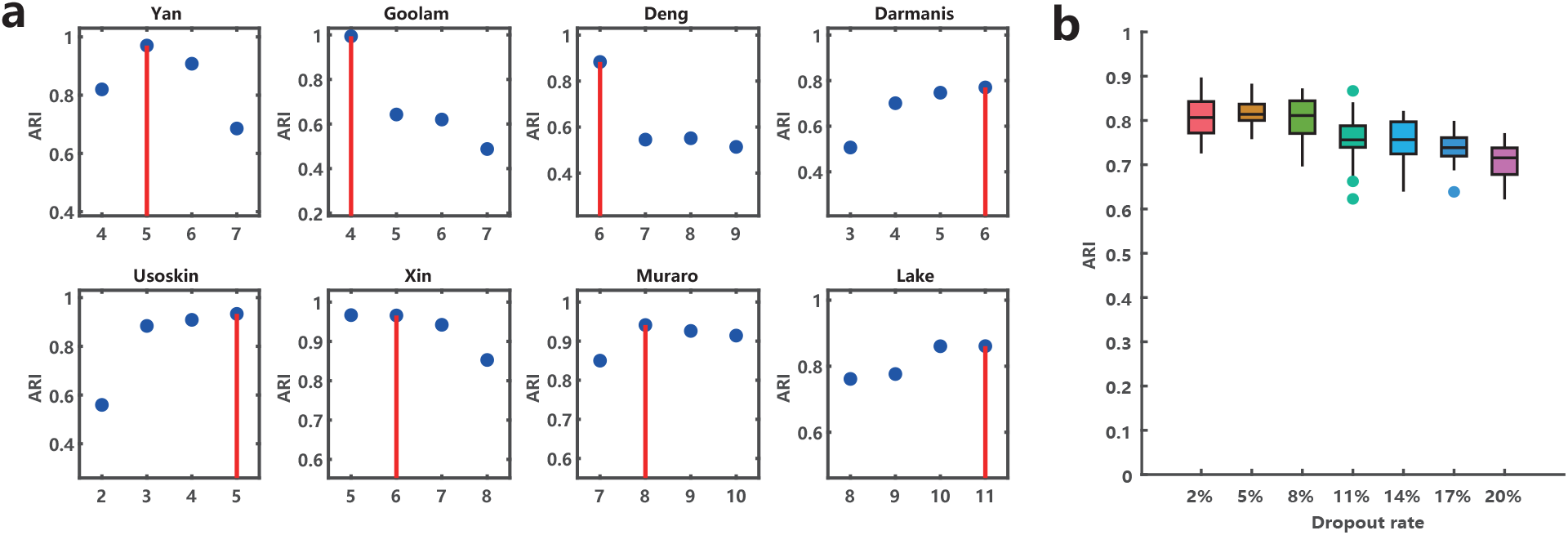
(a) The number of cell types ascertained by AIGS. The red line represents the number of clusters determined by AIGS. (b) Boxplots of clustering accuracy (ARI) after introducing random dropouts ranging from 2% to 20% in eight datasets, with 20 repeated runs.

Despite the robustness of the AIGS framework, occasional discrepancies can arise between the estimated cell type count and the reference number. We have constructed Sankey diagrams to visually represent the relationships between reference cell types and our clusters across all eight datasets (**Supplementary Fig. S8-S15**). There are four principal factors that may contribute to these disparities:

- AIGS has a tendency to cluster cells that display pronounced similarity in functionality or developmental stages, even though they might be acknowledged as distinct types. This tendency was evident in the studies conducted by Yan [12] and Goolam [13].
- If a dataset incorporates a cell class that can be subdivided into multiple sub-types, AIGS has the capacity to discern and segregate these sub-types, as exemplified by the findings of Deng [14] and Usoskin [16].
- In instances where a dataset encompasses a cell class bearing ambiguous characteristics, AIGS tends to allocate these cells into more definitively characterized classes, as was observed in the study by Muraro [18].
- AIGS might not recognize rare cell classes comprising no more than 10 cells, as demonstrated by Xin’s findings [17].

It is noteworthy to mention that the deviations between the number of categories estimated by AIGS and the reference categories predominantly stem from AIGS’s exclusive reliance on gene expression data. For instance, in the analysis of the Yan dataset, AIGS accurately categorized cells from most stages. However, it clustered zygote and 2-cell stage cells into a single class. This particular clustering can be rationalized by understanding the biological processes transpiring during these early developmental stages, particularly the maternal-zygotic transition (MZT), a key event where control over embryonic development transitions from maternal transcripts and proteins to the zygotic genome [20]. Therefore, the gene expression profiles of zygotes and 2-cell stage embryos may bear similarities due to the mutual influence of maternal transcripts and the early stages of zygotic genome activation (ZGA) [21]. This overlapping influence could elucidate why AIGS groups these two stages together, thereby spotlighting the intricacies of cellular clustering based on gene expression data.

### 2.6 Stability on dropout

Missing gene expression, manifesting as an increase in zero and near-zero counts in the dataset, presents a significant challenge in scRNA-seq data analysis. Indeed, in the eight datasets utilized in our paper, the mean dropout rate reached as high as 69% (**Supplementary Table 3**). Despite these constraints, AIGS showcased considerable resilience against such data omissions, as substantiated by the clustering accuracy detailed previously. To further evaluate AIGS’s robustness against dropout phenomena, we conducted an experiment wherein zero-value expressions in the dataset were artificially inflated. Specifically, we randomly assigned a value of 0 to random positions within the dataset, ranging from 2% to 20%. Subsequently, we evaluated the performance of AIGS on these modified datasets. The findings indicate that AIGS demonstrates impressive robustness against dropout events, exhibiting a slow decrease in average clustering accuracy as the quantity of zero-valued expressions expands. Moreover, AIGS’s performance remains relatively steady when dropout is incorporated at various locations, thereby highlighting its robustness in dealing with dropout events (**Fig. 4b** and **Supplementary Fig. S16**).

### 2.7 Visualization and sub-types recognition

In the AIGS pipeline, we have integrated a visualization module to efficaciously delineate the distinctions among cell populations. **Fig. 5a** showcases a comparative display of visual results derived from AIGS, CIDR, scDHA, Seurat, and SIMLR on Yan[12] and Goolam[13] datasets incorporating six stages of embryonic stem cells: zygote, 2-cell, 4-cell, 8-cell, 16-cell, and blast. Additional visualization outcomes can be found in **Supplementary Fig. S17**. This comparative visual analysis underscores the superiority of AIGS in segregating heterogeneous cells and congregating cells of identical types. In order to objectively evaluate the effectiveness of the visualization, we employed the Silhouette Index (SI Score, **Supplementary Note 2.1**) as an evaluative metric. The analysis revealed that AIGS consistently achieves the highest or second-highest SI scores (**Supplementary Fig. S18**). Additionally, the one-sided Wilcoxon signed-rank test manifests a significant enhancement in SI scores for AIGS as compared to other methodologies, with a p-value of 1.1×10^−6^. In datasets such as Yan [12] and Goolam [13], AIGS’s visualization strategypredicated on a sparse-graph representationresults in high silhouette scores due to the intimate within-class connections and sparse between-class connections. This approach engenders clear delineation of inter-class data points in the 2D space while preserving closeness among intra-class data points. In addition to this, compared to commonly used visualization algorithms like t-SNE and UMAP, AIGS’ visualization results show a significant improvement in silhouette scores (**Supplementary Fig. S19** and **20**).

**Figure 5:**
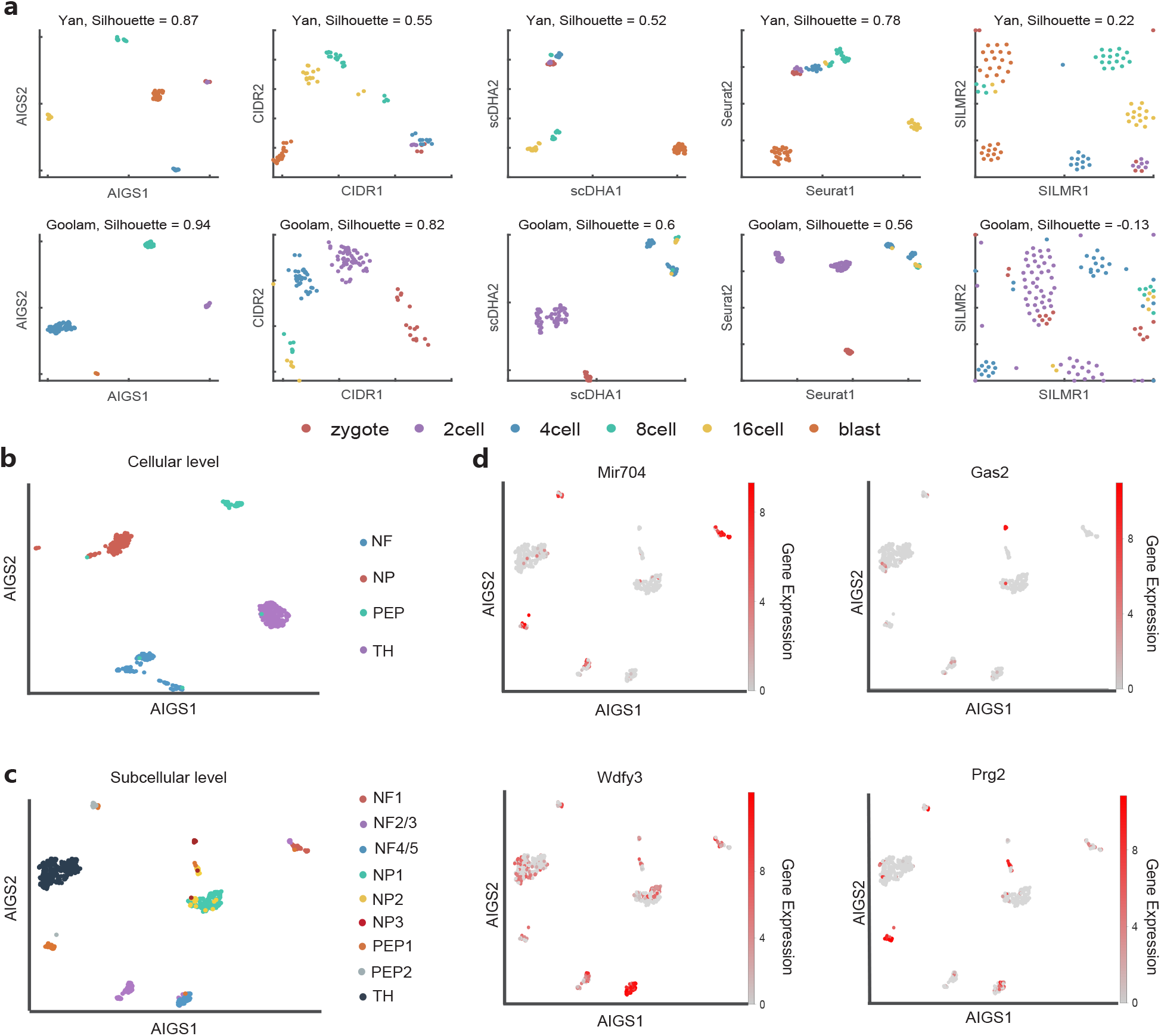
Visualization and marker gene analysis result in cellular level and subcellular level identification. (a) The color-coded visualization result of AIGS, CIDR, scDHA, Seurat, and SIMLR(from left to right) on Yan and Goolam (From Top to Bottom). Each color scatter represents a type of cell. AIGS offers a more accurate reflection of the high-dimensional relative positional relationships of cells, as demonstrated by the higher silhouette coefficients. (b) Visualization results of the Usoskin dataset at low resolution (cellular resolution) under AIGS. (c) Visualization results of the Usoskin dataset at high resolution (Subcellular resolution) under AIGS. (d) The high-resolution visualization results of the Usoskin dataset, with cells colored by the log-normalized gene expression of Mir704, Gas2, Wdr16, and Prg2 in those cells, respectively.

Moreover, AIGS is capable of identifying biologically meaningful cell sub-types and their associated marker genes. Firstly, this capability is demonstrated in our analysis of the Deng dataset [14]. AIGS’s clustering process successfully subdivided these 2-cell stage cells into two distinct sub-types. Upon visualization (**Supplementary Fig. S21a**), these two sub-types appear closely linked, while maintaining their unique identities. This nuanced clustering might signify intricate changes occurring within the 2-cell stage. The marker genes (**Supplementary Fig. S21b**) for these two sub-types might represent two distinct phases of monoallelic gene expression, each contributing to the overall shared expression profile, which is consistent with the findings of Deng et al.[14], who noted that the maternal RNA predominates the early two-cell stage, but gradually diminishes in subsequent stages, reaching parity with paternal RNA at the four-cell stage. The parallel consistent marker gene expression (**Supplementary Fig. S21b**) observed in the 8cell and 16cell stages could denote periods where gene expression dynamics follow a more predictable pattern. It possibly underlines periods of developmental stability that contrast the seemingly chaotic transition seen in the 2-cell stage. In summary, the AIGS application to the Deng dataset not only reinforces the strength of the algorithm in discerning hidden cellular sub-types and their marker genes but also offers fresh insights into the realm of monoallelic gene expression in early cell development stages. Similar analysis results were obtained on the Yan dataset as well (**Supplementary Fig. S22**).

Furthermore, AIGS possesses the capability to discern biologically significant cellular subtypes and pinpoint their associated marker genes by high-resolution visualization. We showcase this proficiency through an analysis of the Usoskin dataset (**Fig. 5b** and **c**), encompassing neuronal fibers (NF), non-peptidergic nociceptors (NP), peptidergic nociceptors (PEP), and cells containing tyrosine hydroxylase (TH). AIGS’s clustering process initially segregates the NF cluster into two discrete subtypes. During low-resolution visualization, both the NF and NP clusters further subdivide into three subtypes each, unveiling close associations within each cluster’s subtypes while preserving their distinctiveness. The fine-grained clustering implies the presence of complex variations within these two clusters, indicating that cells within these clusters exhibit intricate and interconnected heterogeneity.

During high-resolution visualization, the NF cluster is further subdivided into three distinct subtypes, while the PEP cluster is split into two distinct subtypes within each group. Similarly, the NP cluster is categorized into three subtypes. These visualizations of subtypes offer unique insights that are not attainable through alternative analytical methods. AIGS yields subtype identification results in alignment with the discoveries made by Usoskin et al. Their identification of subtype-specific genes, representing expression differences among various subgroups, was accomplished through PCA and outlier identification methods. Additionally, AIGS identifies distinctive marker genes, including Mir704 for the NF1 subtype, Wdfy3 for NF4/5 subtypes, Prg2 for the PEP1 subtype, and Gas2 for the NP3 subtype (**Fig. 5d**). The expression of the Gas2 gene has an impact on cell apoptosis, and its pronounced expression in the NP3 subtype implies that cells within this subtype might be in the concluding stages of NP-type cell development. This observation can influence the subsequent pseudo-time inference process during analysis. Such applications of AIGS lends itself to a deeper understanding of the variability and adaptability inherent in cellular development and the stochastic gene expression processes.

### 2.8 Efficiency of AIGS

In AIGS, the most time-consuming step involve finding the spectral projection, which require the eigenvectors of a symmetric matrix. Since the similarity matrices constructed by AIGS are sparse matrices, we employ the Lanczos[22] algorithm to compute the eigenvectors in our experiments. Both theoretically and experimentally, the Lanczos method has been shown to be superior to other algorithms for solving eigenvector problems in large-scale symmetric sparse matrices. Benefiting from these efficiencies, AIGS’s computational time is under 5 seconds for all datasets, barring the Lake dataset, where it takes 21 seconds. When benchmarked against the leading algorithm, scDHA, AIGS emerges superior by executing seven times faster on the Lake dataset without compromising on accuracy. Additionally, AIGS surpasses scDHA’s operation time by a factor exceeding 20 for other datasets, which underlines its exceptional computational efficiency (**Supplementary Table 4**).

## 3 Discussion

In the realm of single-cell RNA sequencing (scRNA-seq) analysis, we face formidable challenges, including a growing number of cells and genes, technical noise, and substantial dropout rates. To tackle these challenges, we’ve devised an algorithm named AIGS, which leverages interpretable gene selection techniques in its analysis. We tested AIGS on different types of single-cell datasets, including different species (such as mice and humans) and different organs (such as brain and embryo). In contrast to the state-of-the-art methods, AIGS attains remarkable clustering accuracy without necessitating hyperparameter adjustments or relying on deep learning techniques. It consistently achieves an average ARI clustering accuracy exceeding 90%. Additionally, AIGS has higher computational efficiency and stronger robustness when dealing with dropout. Our results show that AIGS generates more accurate visualization results and reliable marker genes, providing a reliable foundation for subsequent biological research.

The success of AIGS can be attributed to two key factors. Firstly, unlike other methods that retain thousands of genes, we have discovered that accurate cell clustering can be achieved by intelligently selecting no more than 300 genes. This significantly enhances analysis efficiency and conserves computational space. Secondly, our newly proposed scale-invariant metric definition remains unaffected by the dimension and density of the data. It is robust enough to handle noise, allowing us to accurately represent the distance relationships between cells. These two advantages together enable AIGS to offer novel biological insights.

In fact, AIGS demonstrates its ability to identify distinct cell subpopulations solely through single-cell sequencing. This is evident in both the clustering and visualization results from the Deng dataset, as well as the varying resolution visualizations from the Usoskin dataset, which align with previously observed biological phenomena. Furthermore, AIGS excels in identifying marker genes corresponding to these subtypes. For instance, the identification of the Gas2 gene as a marker for the NP3 population, characterized by its strong expression, suggests that NP3 may be in the final stages of functional differentiation within the non-peptidergic nociceptors cell population. This finding hints at impending changes in cell fate. This novel biological discovery has the potential to guide the inference of time trajectories for sensory neurons and more.

In conclusion, AIGS has demonstrated to be a powerful tool for scRNA-seq data analysis, capable of accurately selecting a small set of marker genes and applying them to help cluster cells, as well as separating unidentified cell sub-types in the data. Although our experiments were focused on scRNA-seq data, our method is only based on general machine learning algorithms, which means that spatial transcriptomics and multimodal single-cell genomics can also benefit from our approach in the future sequencing and analysis.

## 4 Methods

We use scRNA-seq datasets as examples. AIGS comprises seven elementary steps that take an input gene expression matrix *X* with dimensions *N* × *M*, where *N* represents the number of cells and *M* represents the number of genes.

### 4.1 Gene filtering

The gene filtering process in AIGS comprises two steps. Initially, genes with a maximum absolute expression value across all cells lower than a customizable threshold, denoted as *δ*_1_ (default set to log_2_ 3), are excluded. Subsequently, genes with a variance smaller than a customizable threshold, represented as *δ*_2_ (default set to 1.5), are also discarded. This filter serves the purpose of excluding genes that exhibit minimal expression or variation across cells, as they may not contribute valuable information for clustering. Additionally, gene filtering aids in reducing data dimensionality, enhancing the efficiency of the analysis.

### 4.2 Construction of a sparse similarity graph

A cell-to-cell distance matrix *D* = (*D*_*i j*_) ∈ ℝ^*N*×*N*^ is calculated using either Euclidean or Spearman metric (see **Supplementary Note 1**). The Spearman metric is used as a default setting since it is more robust to gene expression magnitude and more representative of cell differences [23]. For two cells *c*_*i*_ and *c* _*j*_, the symmetric dissimilarity between *c*_*i*_ and *c* _*j*_ is defined as the minimum of two order-based dissimilarities:

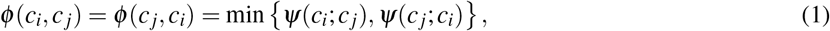

where *ψ*(*c*_*i*_; *c* _*j*_) and *ψ*(*c*_*j*_; *c*_*i*_) provide a scale-invariant dissimilarity through the order of *c* _*j*_ and *c*_*i*_ in each other’s neighbor:

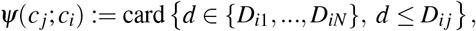

and

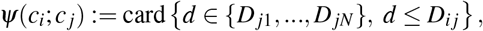

where card{·} denotes the cardinality (number of elements) of a set.

A sparse similarity matrix *S* = (*s*_*i j*_) ∈ ℝ^*N*×*N*^ is constructed using the metric *ϕ* and Gaussian kernel function, defined as:

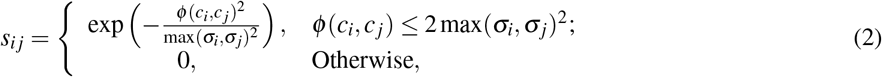

Here, *σ*_*i*_ refers to the *p*-th smallest value among the set {*ϕ*(*c*_*i*_, *c*_1_), …, *ϕ*(*c*_*i*_, *c*_*N*_)} and the value of *s*_*i j*_ indicates the similarity between two cells, *c*_*i*_ and *c* _*j*_, based on their proximity. The parameter *p* represents a confidence level, indicating all cells *c*_*i*_ whose distance from *c*_*i*_ is less than σ_*i*_ and the similarity is at least 0.5. As the distance decreases, the similarity gradually approaches 1. We observed that the majority of cells and their nearest 6 cells are of the same cell type, so *p* is set to 7 to ensure a high confidence level.

### 4.3 Gene selection via learned pseudo-labels

The gene selection step is crucial for identifying cell-type-related genes and reducing noise in single-cell RNA sequencing data. It consists of two stages, as described below:

1. **Learning pseudo-labels**. The learned pseudo-labels 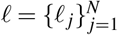 are obtained by the K-means clustering of the spectral projection *H* ∈ ℝ^*M*×*d*^ (**Supplementary Note 3.1** and **3.2**), where *H* is the solution of

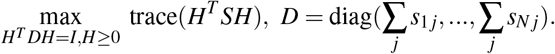

The number of columns *d* in *H* aligns with the pre-clustered classes, and we set it to 3, 4, 5 based on the following considerations: Almost all single-cell RNA datasets can be partitioned into at least three classes. Additionally, we aim to avoid an excessive number of pre-clustered classes, as this could lead to the fragmentation of certain cell types, thereby impacting subsequent gene selection processes. It is important to note that K-means is a greedy algorithm, which may produce different locally optimal solutions based on different initial values. Therefore, we randomly repeat K-means 20 times for each clustering and automatically select the optimal result using the objective function of K-means as a criterion.
2. **Selecting cell-type-related genes**. In an expression matrix, each gene can be represented as a column vector of length *N* denoted by *g*_*i*_. AIGS selects genes that are consistent with the learned pseudo-labels 𝓁 ∈{𝓁^(3)^, 𝓁^(4)^, 𝓁^(5)^}, where the consistency between each *g*_*i*_ and 𝓁 is measured by the normalized mutual information (NMI) between the cluster labels 𝓁 and the expression profile of gene *g*_*i*_. To this end, we quantize *g*_*i*_ as

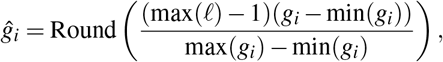

where the function Round(·) in *ĝ*_*i*_ rounds a number to its closest integer. Hence, the range of *ĝ*_*i*_ is equal to that of 𝓁 and we are able to compute the NMI (**Supplementary Note 4**) between *ĝ*_*i*_ and 𝓁. For each pesedo-label 𝓁, we select *k* (default as 100) genes that achieve the highest NMI. The final selected genes consist of a union of the gene groups identified by 𝓁^(3)^, 𝓁^(4)^, and 𝓁^(5)^. With these selected genes, we represent each cell as a vector with much less dimension than that in the original dataset, denoted by *ĉ*_*j*_. By selecting these cell-type-related genes, we can provide more accurate descriptions of the features of each cell, and reduce the impact of noise and redundant information, thereby improving the accuracy of analyzing and interpreting single-cell RNA sequencing data.

### 4.4 Removing Outlier

Due to technical noise arising from amplification biases, batch effects, and sequencing errors, the readings of some cells may have been out of calibration, resulting in some outliers in *ĉ*_*i*_. These outliers may contain some doublets or empty drops, which might interfere with clustering. To address this issue, AIGS provides an outlier detection module to detect and remove outliers. With the new cell representation *ĉ*_1_, …, *ĉ*_*N*_ obtained from selected genes, we recalculate a Spearman distance matrix 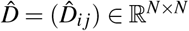, and define the aggregation degree **Agg** for each *ĉ*_*i*_ as:

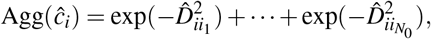

where 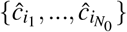 are the *N*_0_ cells closest to *ĉ*_*i*_. A smaller Agg(*ĉ*_*i*_) implies that *ĉ*_*i*_ is far away from its neighbors and therefore more likely to be an outlier. AIGS selects η% (5% as default) of all cells with the smallest Agg(*ĉ*_*i*_) and removes them from the dataset. This process helps to improve the accuracy of clustering by removing outlier cells that might interfere with the clustering results.

### 4.5 Estimate the number of cell types and Clustering

Let 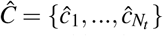 denote the dataset without outliers, where *N*_*t*_ is the number of remaining cells and each cell *ĉ*_*i*_ is represented by the expression value of the selected genes. We construct a Boole matrix 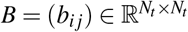 as follows.

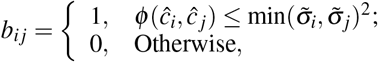

where *ϕ*(·,·) is defined as (1) and 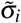 refers to the third smallest value among the set 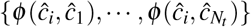. Cells of the same type tend to have similar gene expression profiles, so they are more likely to be connected to each other in B, forming a connected subset. Thus, we compute the number of connected components of *B* as an initial estimate of the number of cell types, denoted as *nc*. The final estimation of the cell type number is chosen from 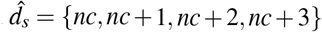. To find the optimal estimate, we construct the similarity matrix *Ŝ* from *Ĉ* as (2) and execute spectral clustering to *Ŝ*, where the number of clusters is selected in 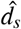. The obtained labels 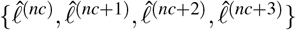 are used to construct the label-graphs *B*^*q*^ as

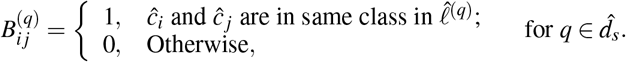

Finally, the number of clusters (cell types) is chosen as the maximum of constituency between the similarity matrix *Ŝ* and the label-graph *B*^(*q*)^

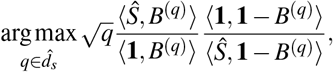

where 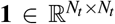 means a all-one matrix and ⟨·,·⟩represents the Frobenius inner product of two matrices (**Supplementary Note 1**).

### 4.6 Clustering and 2D Visualization

To derive the final cell type labels 𝓁, we utilize the spectral projection 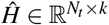 from similarity matrix *Ŝ*, where *k* is the estimated cell-type number, and perform K-means clustering. For visualizing the cell types and their relationships, we have adopted a visualization framework similar to the UMAP algorithm [24], recognizing the critical importance of initialization and graph construction in the effectiveness of the UMAP algorithm[25]. Therefore, we have enhanced its initialization and graph construction components while preserving the optimization module for the loss function. Initially, we leverage the outstanding clustering results of AIGS to provide a robust visualization.

Let the cell division given by the clustering result of AIGS be {*C*_1_, · · ·, *C*_*k*_} and the spectral projection result be 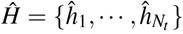. First, the cluster center *c*_*i*_ of *C*_*i*_ is calculated, and the normalized Euclidean distance *D*_*i*_ = *D*_*i*_(*j*) from each sample in *C*_*i*_ cluster to its corresponding cluster center is calculated as follows:

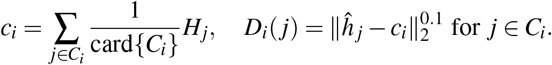

In 2D, we distribute the cluster center on the unit circumference as 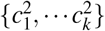, that is, the 2D coordinate of the cluster center is

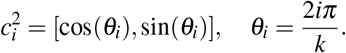

Finally, we construct the initial embedding of *H* in 2D is

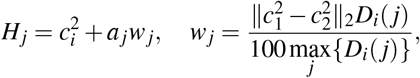

where *a*_*i*_ is a random vector generated according to standard two-dimensional normal distribution.

In terms of graph construction, we also employ scale-invariant distances between cells. Drawing inspiration from the ResNet concept, we utilize clustering results to further enhance the similarity between cells of the same type. AIGS first standardizes the spectral projection as follows:

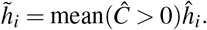

Here, *Ĉ >* 0 refers to selecting all elements in the matrix that are greater than 0 to form a vector, and the mean value of this vector is calculated using the function mean(·). After this, AIGS proceeds to reconstruct the samples 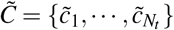 as follows:

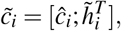

where 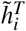 means the transpose of a vector 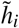.

Finally, we reconstruct the similarity matrix 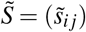 as the input visualization as

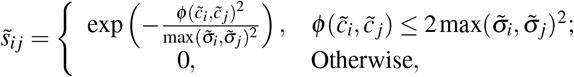

where 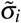 is defined the same way as in the original graph setting. The spectral projection is a valuable tool that increases intra-class similarity and decreases inter-class similarity compared to the original data. Consequently, by integrating the spectral projection with the original data, the reconstructed similarity matrix can more accurately portray the actual distribution of cells in high-dimensional space. Therefore, AIGS can produce more precise visualizations with the assistance of this improved similarity matrix.

Consider the minimization problem within the UMAP framework,

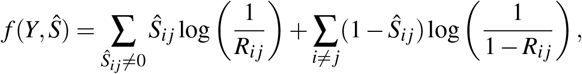

where 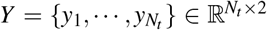 is the 2D embedding of cells, and 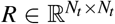 is the similarity matrix of *Y*. The similarity between two points is defined as

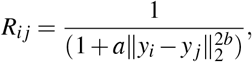

where *a* and *b* are constants and further information on the selection of hyper-parameters, as well as the iterative steps, can be found in [24] and **Supplementary Note 5**.

For the visualization at the cellular subtype level, our objective is to minimize the similarity between different cell subtypes as much as possible. Therefore, we further sparsify the similarity matrix 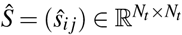 as:

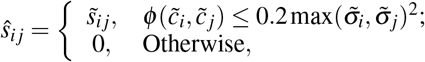

Similarly, by minimizing *f* (*Y, Ŝ*), we obtain a visualization result *Y* that enhances the resolution of cellular subtypes. Our method can more accurately reflect the distribution of cells in high dimensions in two dimensions compared to existing excellent visualization methods, such as t-SNE[26] and UMAP[24] (**Supplementary Fig. S19** and **S20**).

### 4.7 Marker gene selection

In the final step of AIGS, we employ the ANOVA test [27] to identify marker genes from the gene expression matrix after gene filtering. This is achieved by comparing the gene expression of different clusters using the clustering results obtained from AIGS.

To perform this test, we start by selecting a gene and assigning one of the cell types in the clustering results as 1, with the rest designated as 0, represented as label 𝓁_0_. We then calculate the means and variances of the gene expression in the 0 and 1 cell types as *m*_0_, *m*_1_ and *v*_0_, *v*_1_, respectively.

Subsequently, we compute the p-value of the ANOVA for the gene expression and the label 𝓁_0_ and set the selection criteria as follows:

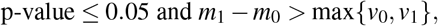

If a gene satisfies the selection criteria, we consider it a marker gene and record the p-value.

Finally, we select the genes with the smallest *k* (default as 30) p-values as the final marker genes. These marker genes offer valuable insights into the specific cell types and their gene expression patterns, allowing researchers to understand the underlying biology of AIGS and develop new therapies (**Supplementary Table 5**).

To further refine the identification of marker genes for cellular subtypes, we begin by re-clustering the visualization results *Y* at a specific resolution level corresponding to our target subtypes. Subsequently, we leverage the clustering results with the proposed marker gene identification steps to complete the marker gene recognition process.

First, we create the Euclidean distance matrix *D*, denoted as 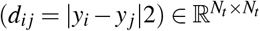, for the visualization results *Y*. Additionally, we generate a Boolean matrix *B*, represented as 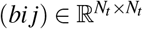, using the following definition:

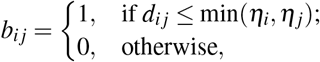

where *η*_*i*_ corresponds to the tenth smallest value among the set 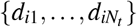.

To identify distinct cellular subtypes, we analyze the connected components of the Boolean matrix *B*. It’s important to highlight that in constructing this Boolean matrix, we employ a 10-nearest neighborhood criterion based on the visualization results. This criterion ensures that cellular subtypes contain a sufficient number of cells, which enhances the robustness of subtype identification. Finally, we utilize the marker gene selection step to obtain marker genes for cell subtypes based on the high-resolution clustering results.

## Supporting information

Supplementary explanations and supporting evidence for the conclusions in the main text of the paper

## 6 Data Availability

The datasets supporting the conclusions of this article are publicly available in the Gene Expression Omnibus (GEO) or other data platforms. They can be found under the following project accession numbers: GSE36552[12], EMTAB3321[13], GSE45719[14], GSE67835 [15], GSE59739 [16], GSE81608 [17], GSE85241 [18], and PHS000833 [19]. The detailed information for each dataset is discussed in the **Supplementary Note 6**. For each mentioned dataset, a logarithmic transformation (base 2) was applied to the raw expression data from individual cells before analysis.

## 7 Availability of Source Code and Requirements

Project name: AIGS.

Project homepage: https://github.com/BioCom-Lab/AIGS.git.

Operating system(s): Windows 10.

Programming language: Matlab and Python.

Other requirements: Matlab >= 2021a; Python>=3.11.5; Numpy>=1.24.3; NetworkX >= 3.1; Scipy>=1.11.1; Scikit-learn>=1.3.0; Matplotlib>=3.8.2.

License: GPL-3.0 license.

RRID: SCR_001622 (Matlab), SCR_008394 (Python).

Remark: Due to the significant number of matrix operations involved, we suggest using the MATLAB version for faster clustering and visualization.

## 8 List of abbreviations

scRNA-seq: Single Cell RNA Sequencing
AIGS: Analyzer with Intelligent Gene Selection
NMI: Normalizaed Mutual Information
ARI: Adjusted Rand Index
MZT: Maternal-Zygotic Transition
ZGA: Zygotic Genome Activation
SI: Silhouette Index
NF: Encompassing Neuronal Fibers
NP: Non-Peptidergic Nociceptors
PEP: Peptidergic Noci-ceptors
TH: Cells Containing Tyrosine Hydroxylase
ACC: Accuracy
ARI: Adjusted Rand Index
Jaccard: Jaccard Coefficient
GEO: Gene Expression Omnibus.

## 9 Ethics Approval

The experiments in this article do not involve human or animal participants.

## 10 Acknowledgements and Funding

This research was supported in part by the Zhejiang Provincial Natural Science Foundation of China under Grant No. LQ22C090007; in part by the National Natural Science Foundation of China under Grant No. T2241018; in part by the National Science and Technology Major Project of the Ministry of Science and Technology of China under Grant No. 2021ZD0114303; in part by the Key Research Project of Zhejiang Lab under Grant No. 2020KB0AC01; in part by National Nature Science Foundation of China under Grant No. 62101390.

## 11 Author contributions

Tianhao Ni, Xinyu Zhang and Kaixiu Jin: Conceptualization (lead); Formal Analysis (lead); Methodology (lead). These three authors made equal contributions to the paper. Guanxiong Pei and Nan Xue: Investigation (lead); Writing-Original Draft Preparation (equal). Guanao Yao: Methodology (equal). Bingjie Li and Taihao Li: Funding Acquisition (supporting), Supervision (lead), Writing - Review and Editing (lead).

