## Supplementary explanations and supporting evidence for the conclusions in the main text of the paper for "Interpretable scRNA-seq Analysis with Intelligent Gene Selection"

### Supplementary Information

#### Contents

**A Supplementary Figures** **2**

**B Supplementary Table** **24**

**C Supplementary Note 1: Definition of distance mentioned** **31**

**D Supplementary Note 2: Performance Metrics** **32**

**H Supplementary Note 6: Data sources of single-cell RNA-seq** **37**

### A Supplementary Figures

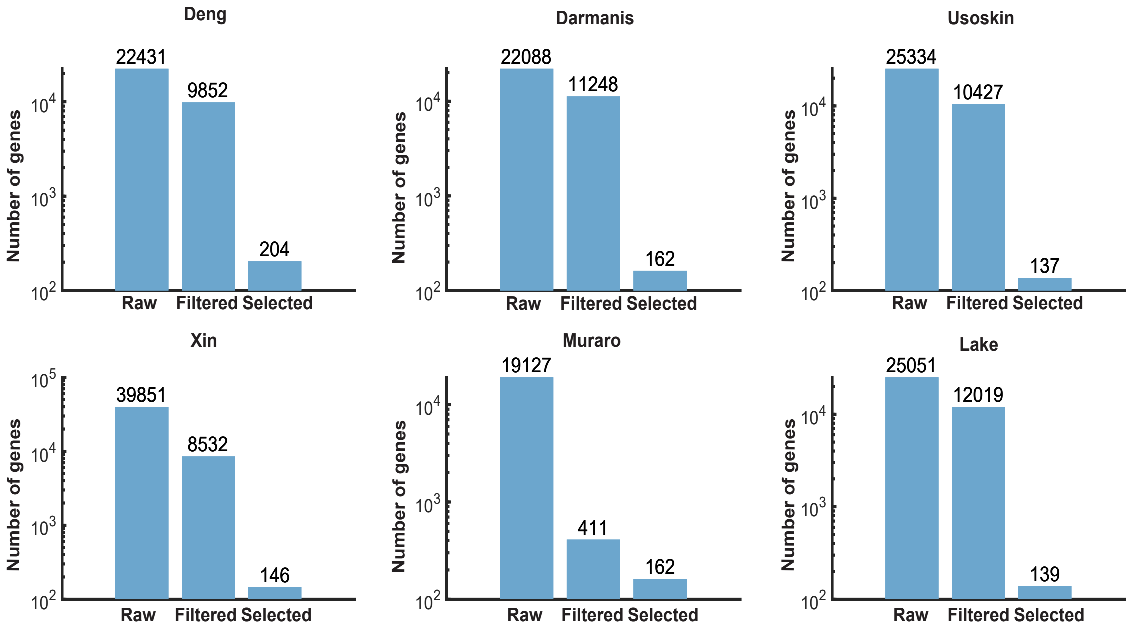

Figure S1: Selected genes during gene filtering and selection in AIGS across six datasets [8, 9, 10, 11, 12, 13]: In each box plot, the first column represents the number of genes in the dataset, the second column shows the remaining genes after filtering, and the third column indicates the remaining genes after gene selection. The algorithm retains only a small subset of genes that contribute to accurate clustering due to the removal of noise from the data.

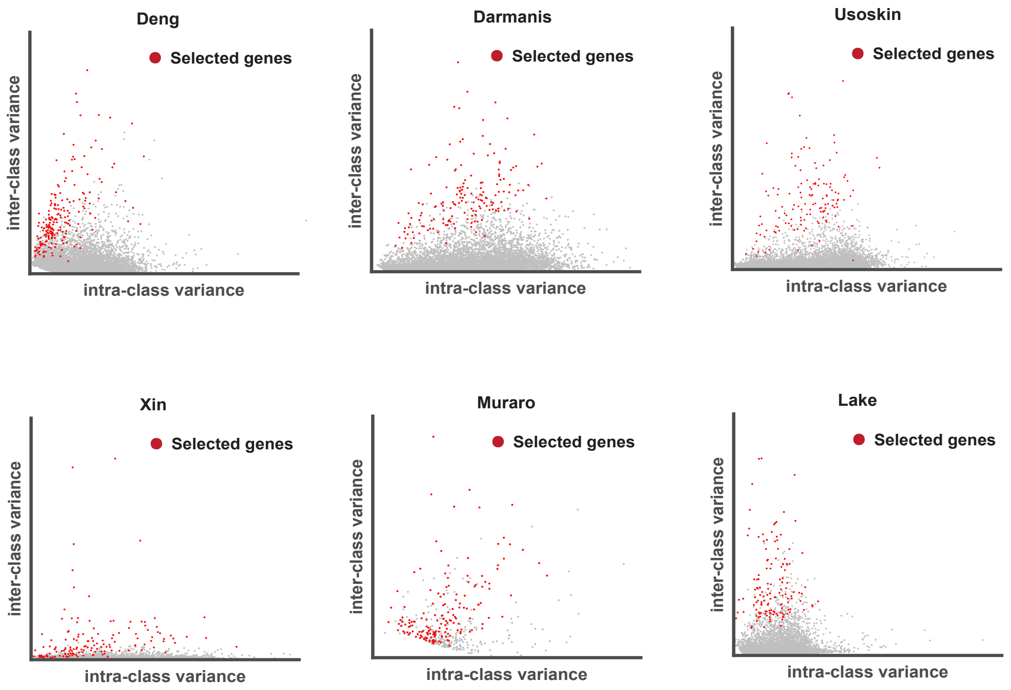

Figure S2: Comparison of selected and eliminated genes intra-class and inter-class variances in six datasets [8, 9, 10, 11, 12, 13]. The red points represent genes selected by AIGSs gene selection module, while the gray points represent genes that were not selected

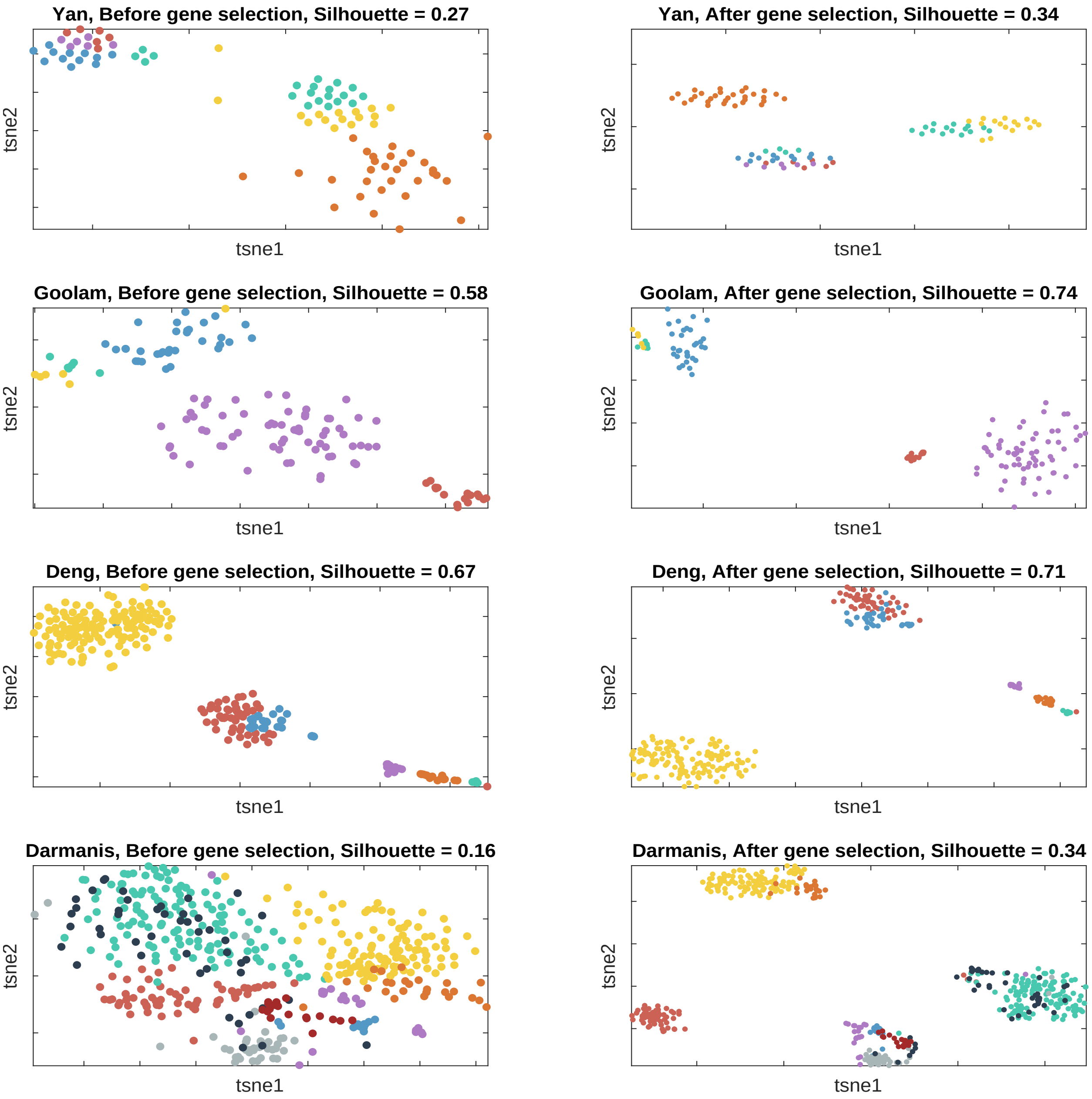

Figure S3: Comparing silhouette coefficients (see **Note D.1**) of t-SNE visualizations [28] before and after gene selection in AIGS on Yan [6], Goolam [7], Deng [8], and Darmanis [9] datasets: The first columns display results prior to gene selection, while the second columns exhibit results following gene selection for the respective datasets. Despite the removal of a large number of genes through the gene selection process, the retained genes continue to produce low-dimensional representations with enhanced silhouette coefficients.

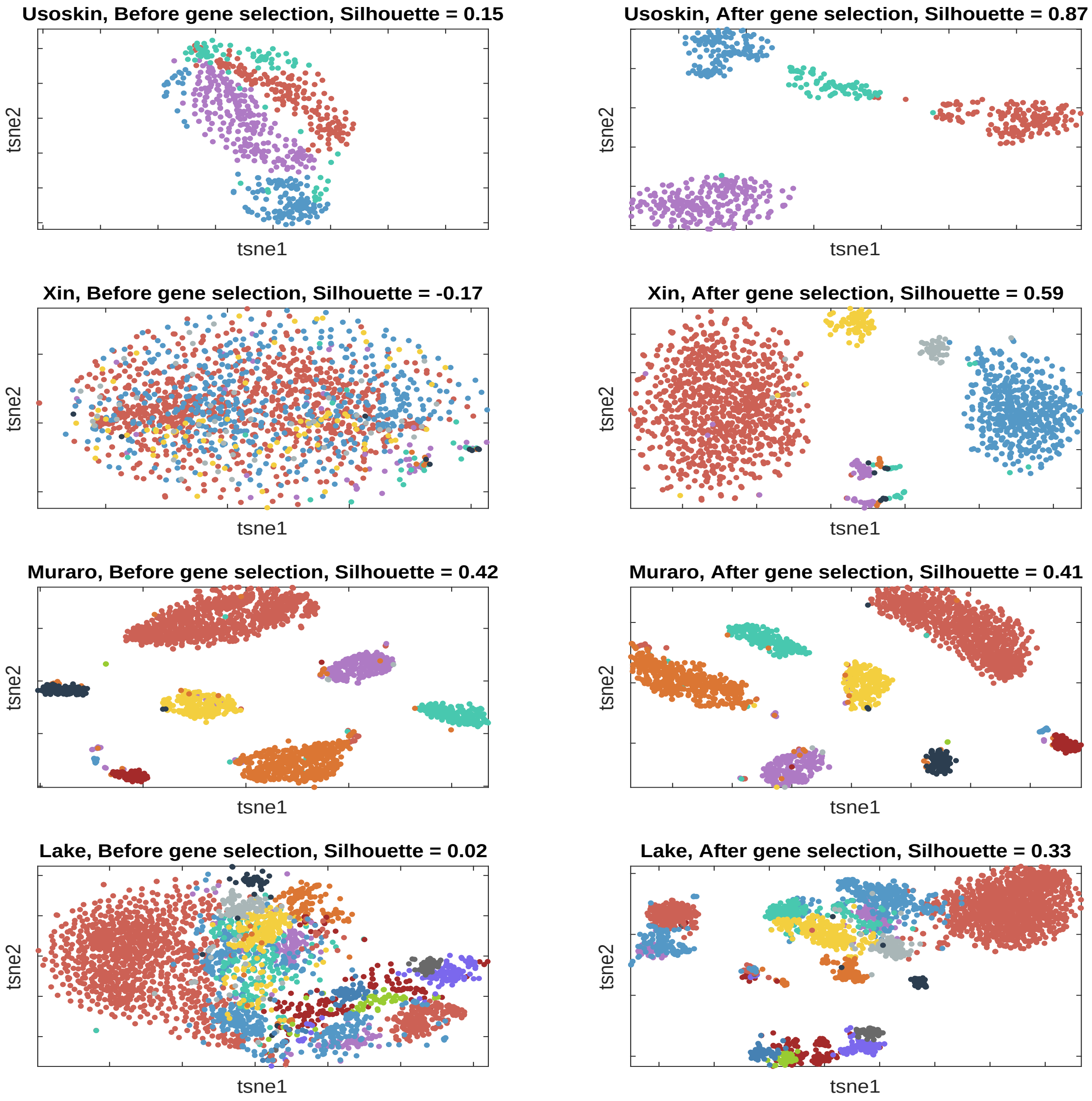

Figure S4: Comparison of silhouette coefficients (see **Note D.1**) of t-SNE visualization [4] before and after gene selection in AIGS on Usoskin [10], Xin [11], Muraro [12] and Lake [13] datasets. The first columns display results prior to gene selection, while the second columns exhibit results following gene selection for the respective datasets.

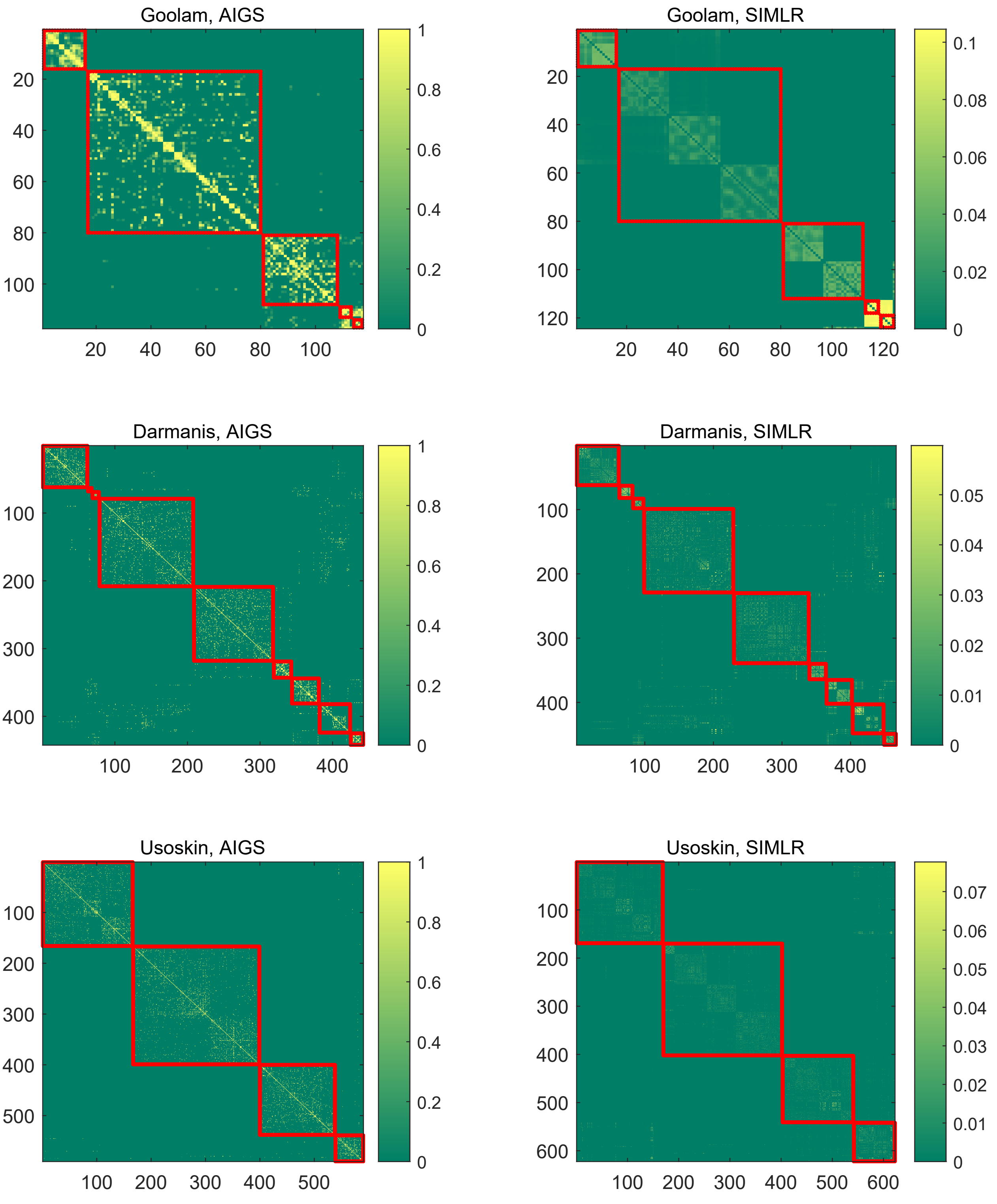

Figure S5: Similarity matrix comparison of AIGS and SIMLR [4] on Goolam [7], Darmanis[9] and Usoskin[10] datasets. Red squares represent the similarity of cells of the same type, while contrasting color boxes represent the similarity of cells of different types. Lighter colors indicate higher similarity. AIGS's scale-invariant graph demonstrates higher intra-class similarity and lower inter-class similarity compared to SIMLR's similarity graph, more accurately representing proximity and separation of cells in high-dimensional space.

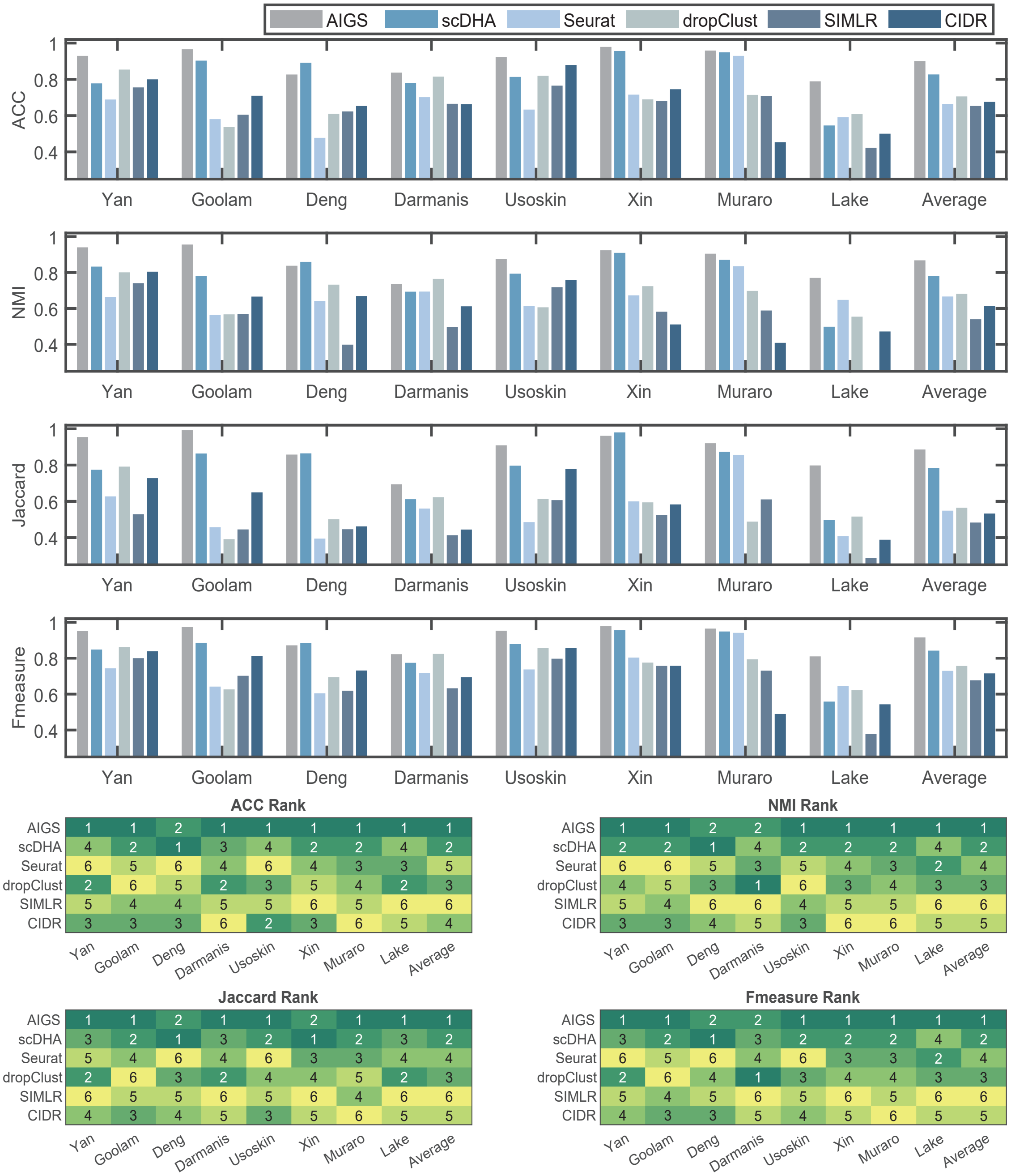

Figure S6: Evaluation of AIGS, scDHA [1], SEURAT [2], DropClust [3], SIMLR [4], and CIDR [5] for single-cell clustering: A performance and ranking comparisons based on Accuracy(ACC), Normalized Mutual Information(NMI), Jaccard Coefficient(Jaccard), and Fmeasure (see **Supplementary Note 4**) across eight datasets [6, 7, 8, 9, 10, 11, 12, 13], with AIGS achieving the highest average clustering accuracy and ranking second in two datasets.

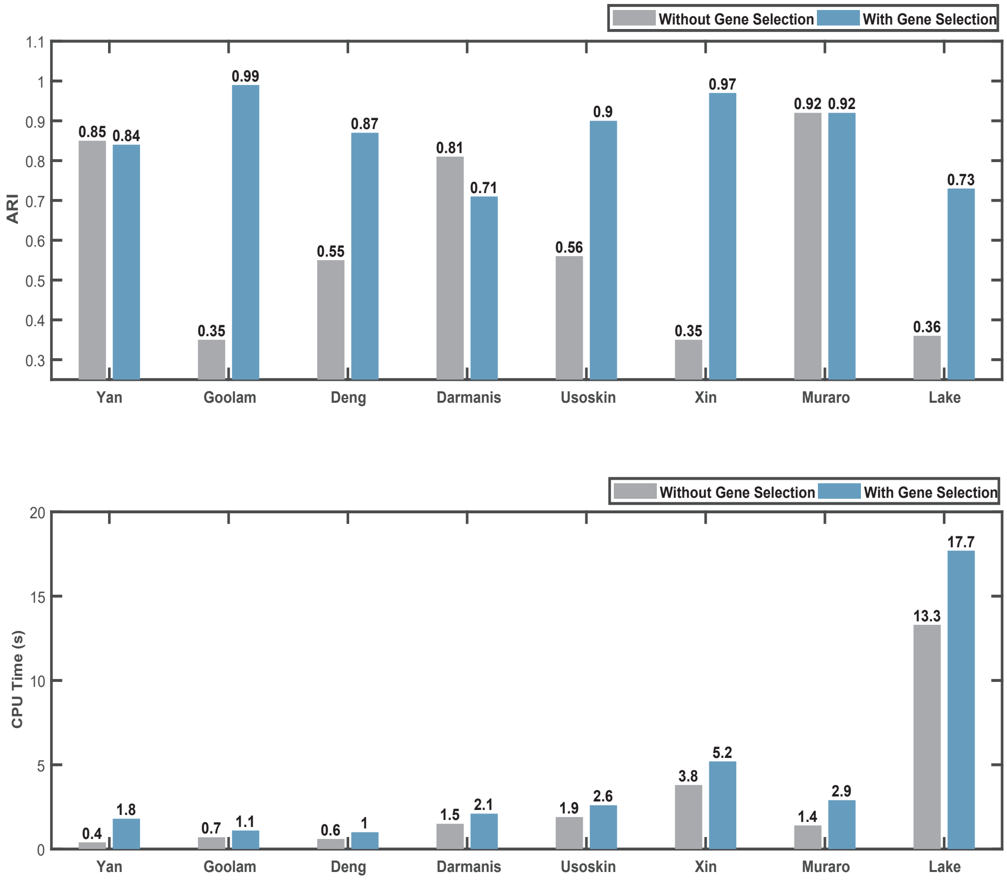

Figure S7: Impact of gene selection on ARI and CPU Time in AIGS clustering: The upper figure displays the comparison of clustering accuracy ARI before and after gene selection across eight datasets [6, 7, 8, 9, 10, 11, 12, 13]. In each dataset, the first column represents the ARI prior to gene selection, while the second column shows the ARI post gene selection. The lower figure illustrates the comparison of CPU time for the clustering step before and after gene selection across eight datasets. In each dataset, the first column denotes the CPU time before gene selection, and the second column highlights the CPU time after gene selection. Gene selection results in a modest increase in CPU time but significantly enhances ARI, demonstrating the effectiveness and efficiency of the approach.

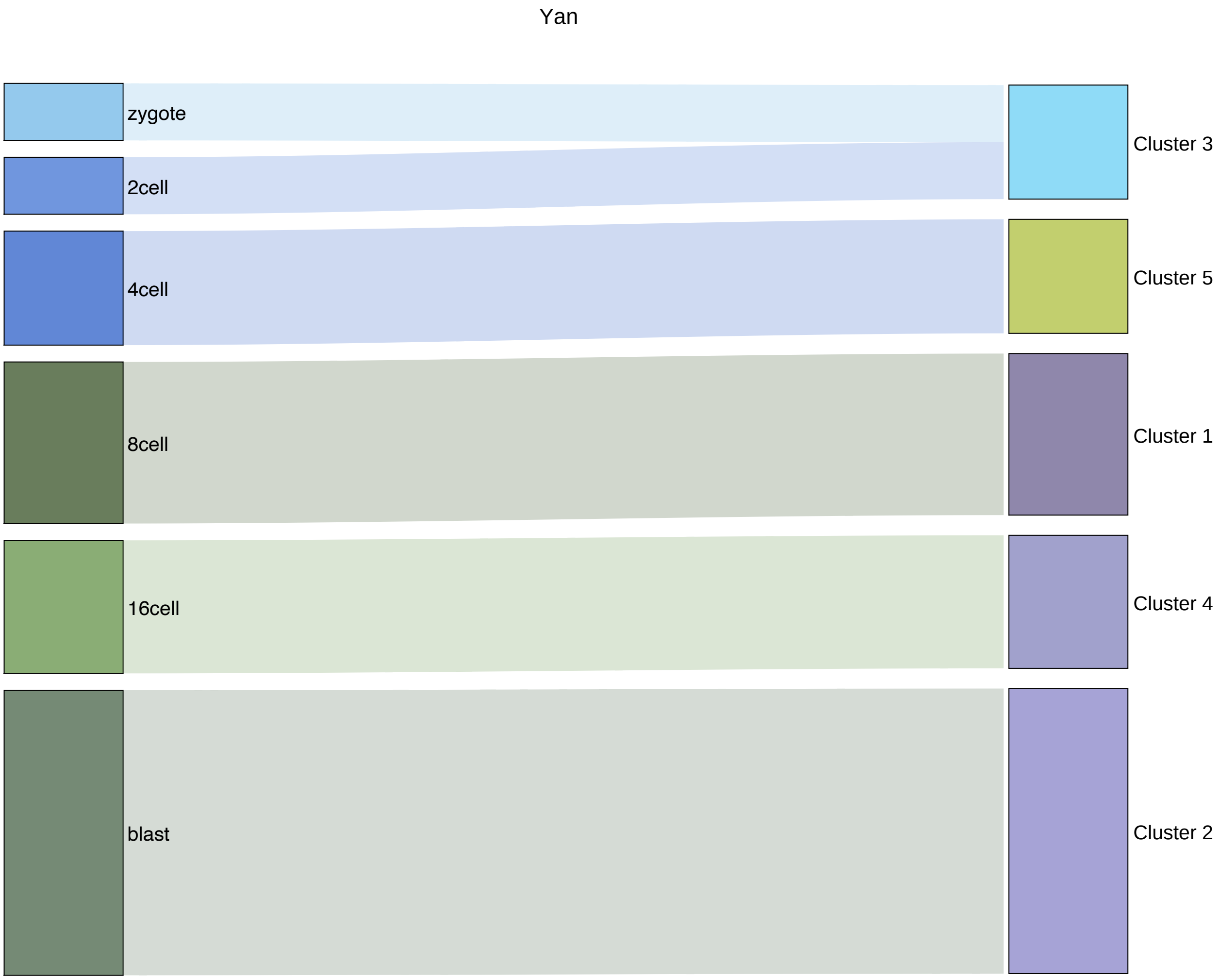

Figure S8: A comparative view of AIGS clustering outcomes with reference labels using a Sankey diagram for the Yan dataset [6]: The diagram's left side showcases the genuine cell classification labels, while the right side provides a visual representation of AIGS's clustering results. Each rectangle symbolizes a cell class, and its length is indicative of the number of cells encapsulated within that class. When applied to the Yan dataset, AIGS amalgamated zygote and 2-cell stage cells into a single category. Nonetheless, it demonstrated precision in clustering cells from the remaining stages, including the 4-cell, 8-cell, 16-cell, and blast stages, into their corresponding categories.

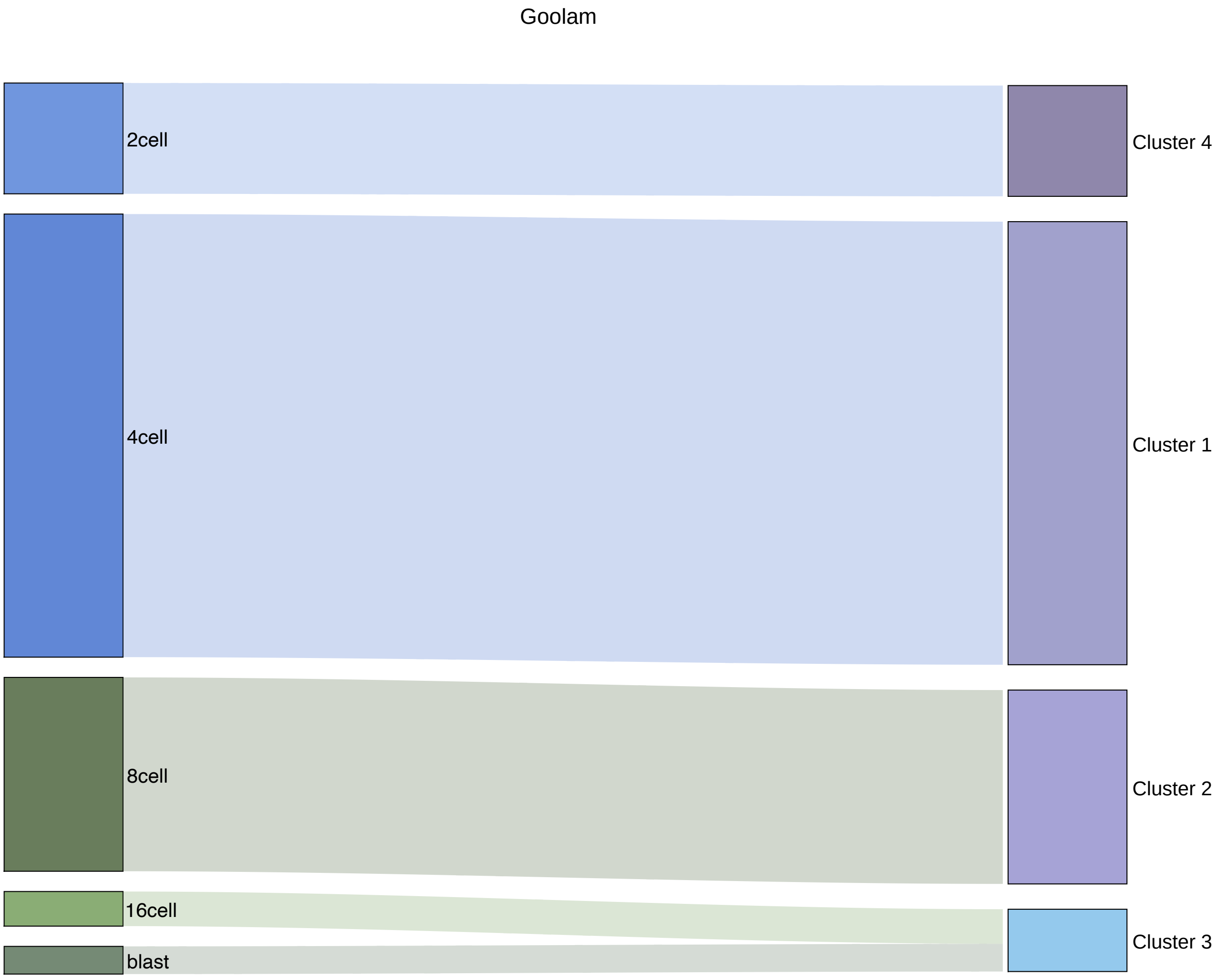

Figure S9: Sankey diagram comparison of AIGS clustering results with gold standard labels on Goolam [7] dataset. AIGS grouped 16-cell and blast stage cells together as one category, while accurately clustering cells from the other stages, including 2-cell, 4-cell, and 8-cell stages, into their respective categories.

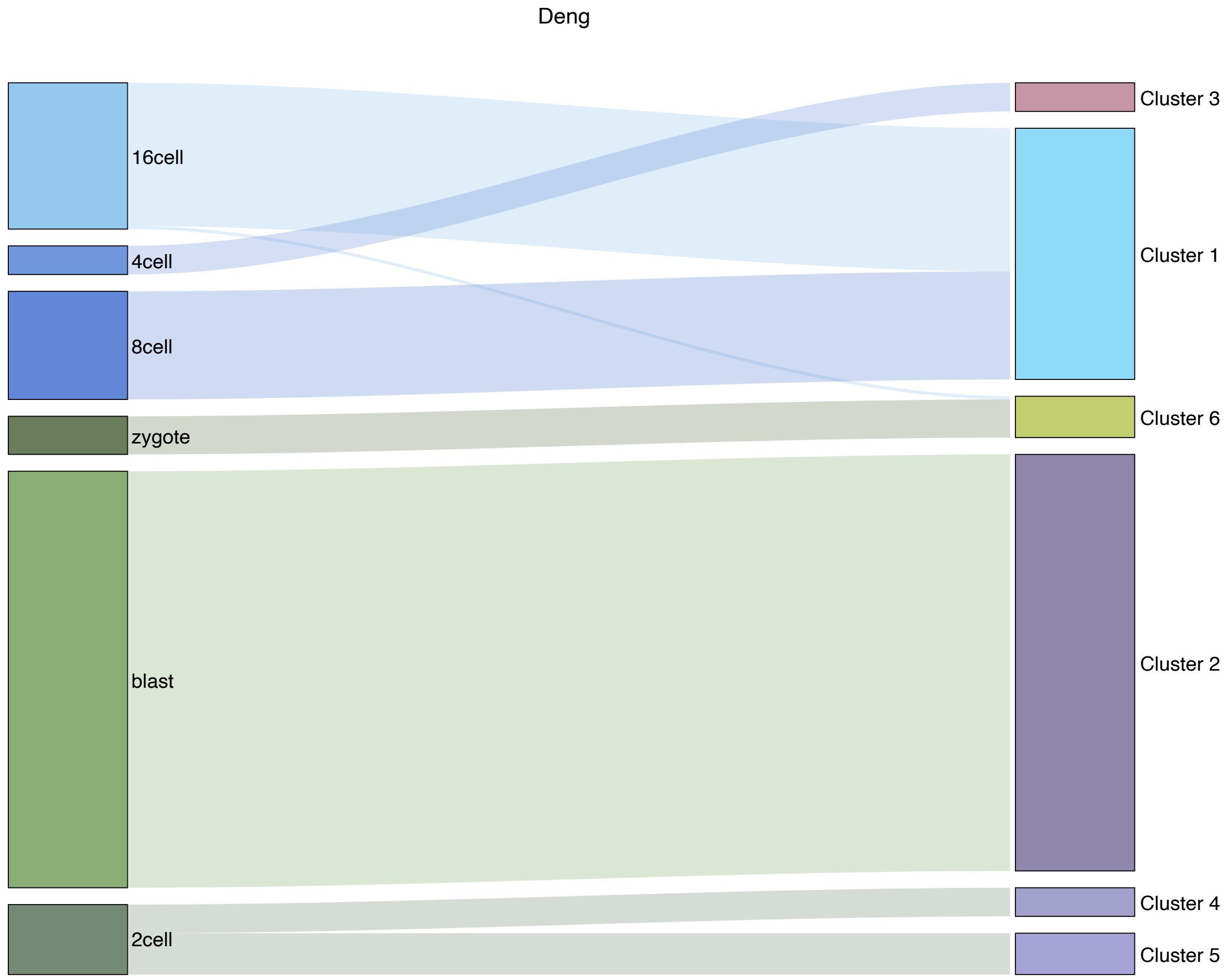

Figure S10: Sankey diagram comparison of AIGS clustering results with gold standard labels on Deng [8] dataset. AIGS grouped 8-cell and 16-cell stage cells together as one category, while considering 2-cell stage cells as two subtypes. Among the remaining cell types, except for grouping one 16-cell stage cell with the zygote stage, AIGS accurately clustered cells from all other stages, including zygote, 4-cell, and blast stages, into their respective categories.

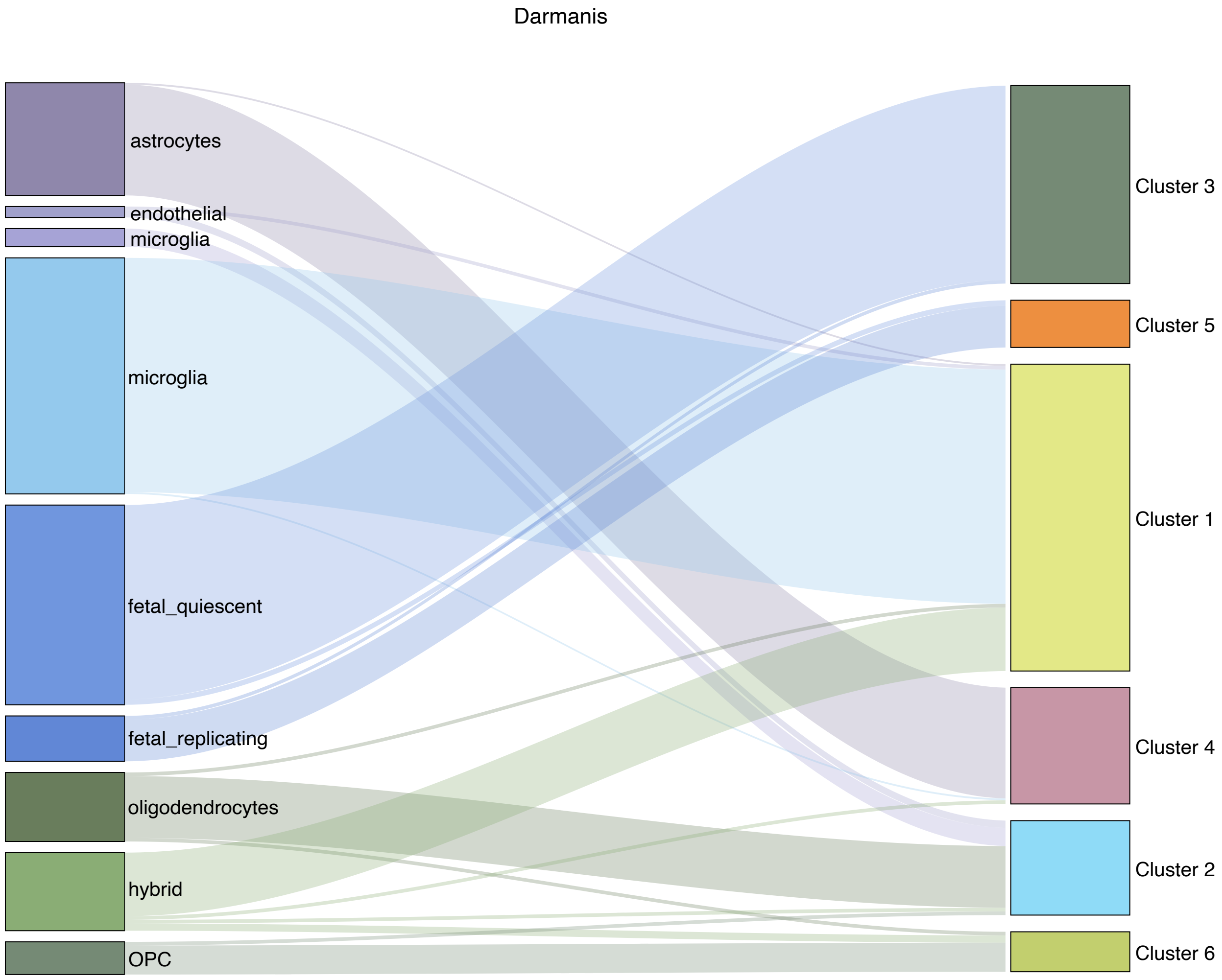

Figure S11: Sankey diagram comparison of AIGS clustering results with gold standard labels on Darmanis [9] dataset. AIGS clustered endothelial cells, microglia, and oligodendrocytes into one category, while grouping microglia and hybrid cells into another category. The remaining cells, including astrocytes, fetal-quiescent, fetal-replicating, and OPC, were mainly clustered into their own categories.

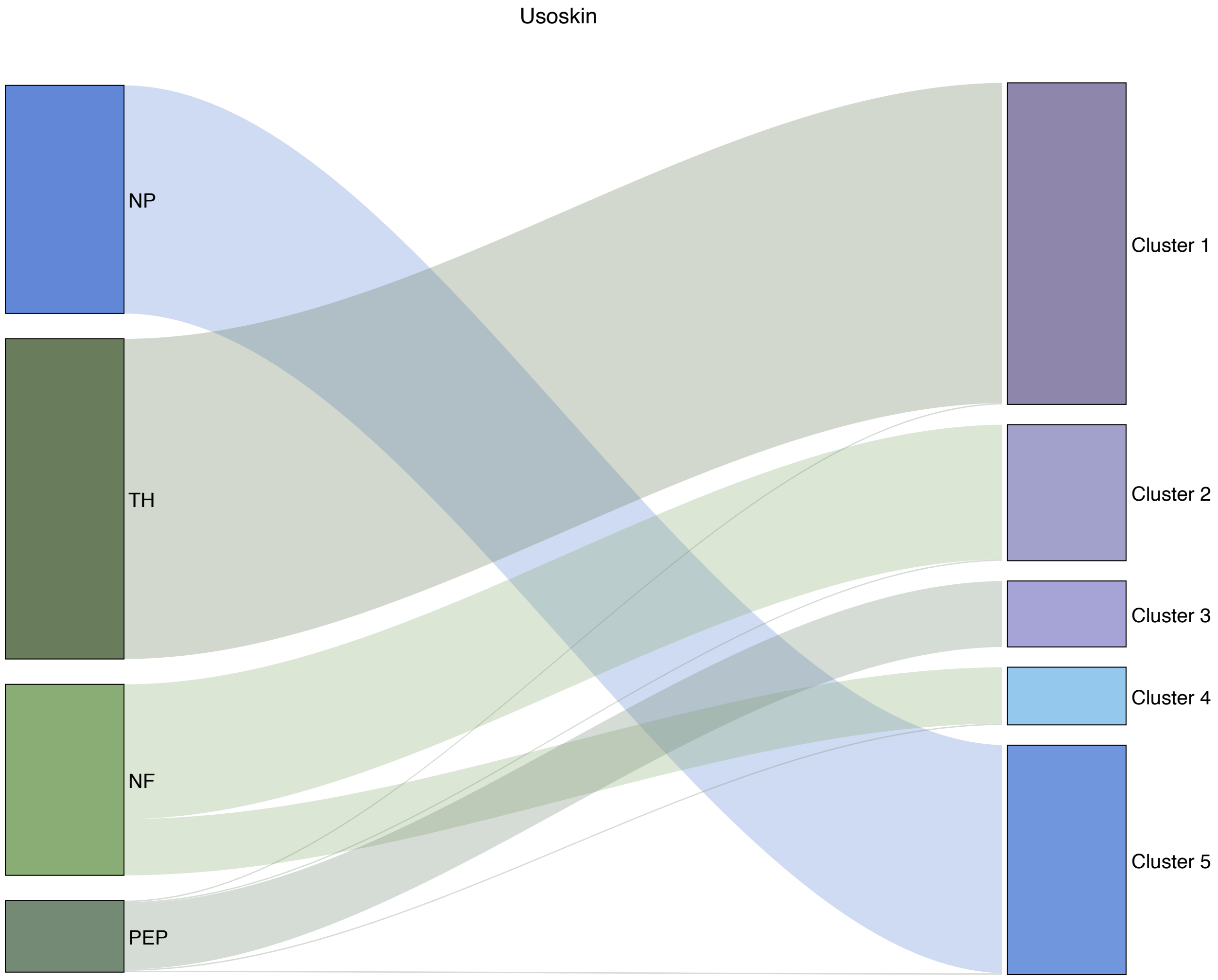

Figure S12: Sankey diagram comparison of AIGS clustering results to golden labels on Usoskin [10] dataset. Besides AIGS's classification of NF into two subtypes, the remaining categories are predominantly classified accurately.

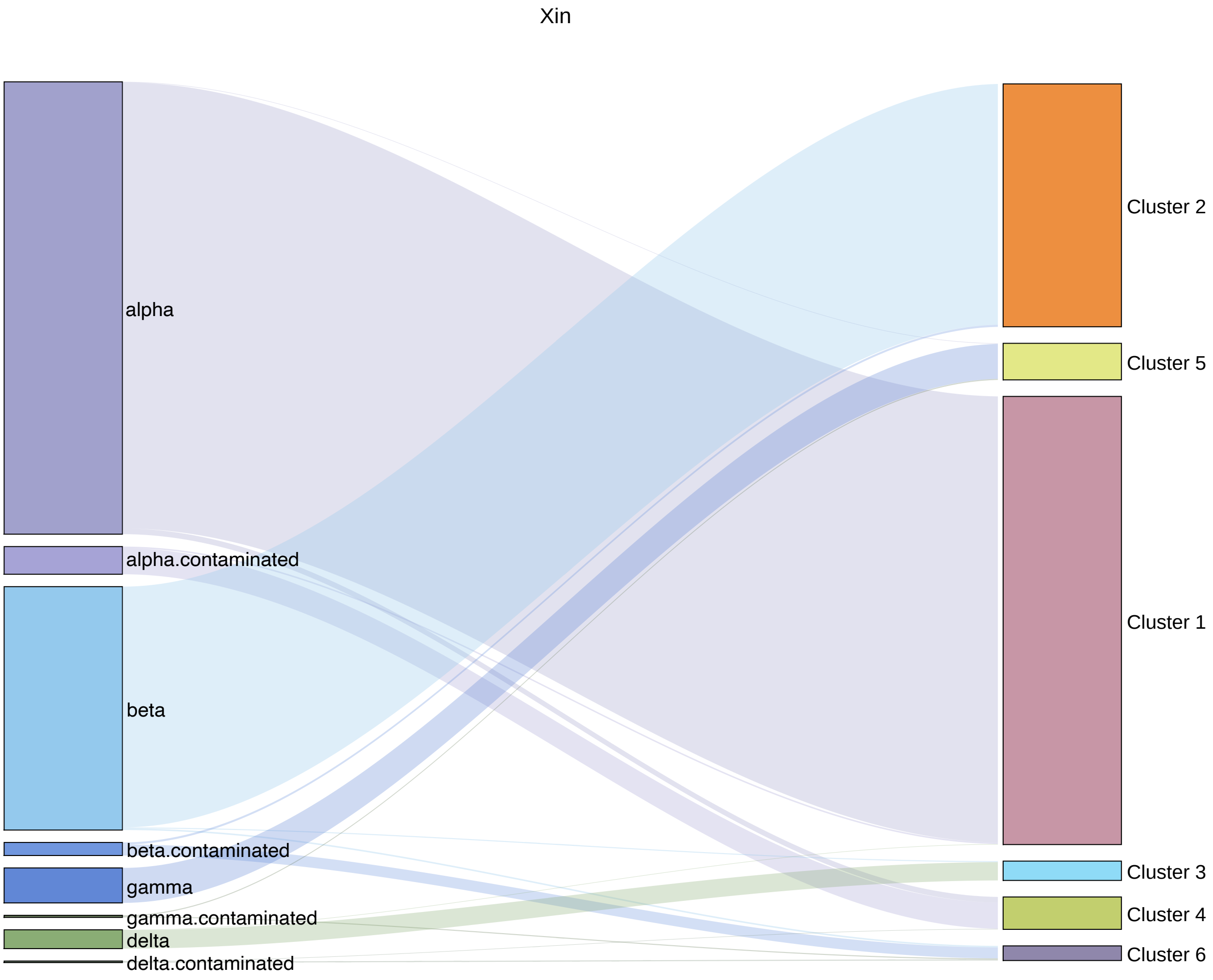

Figure S13: Sankey diagram comparison of AIGS clustering results with golden labels on Xin [11] dataset. AIGS placed the gamma-contaminated and delta-contaminated cell types with very few cells into the other category. The alpha, alpha-contaminated, beta, beta-contaminated, gamma, and delta cell types were accurately classified by AIGS.

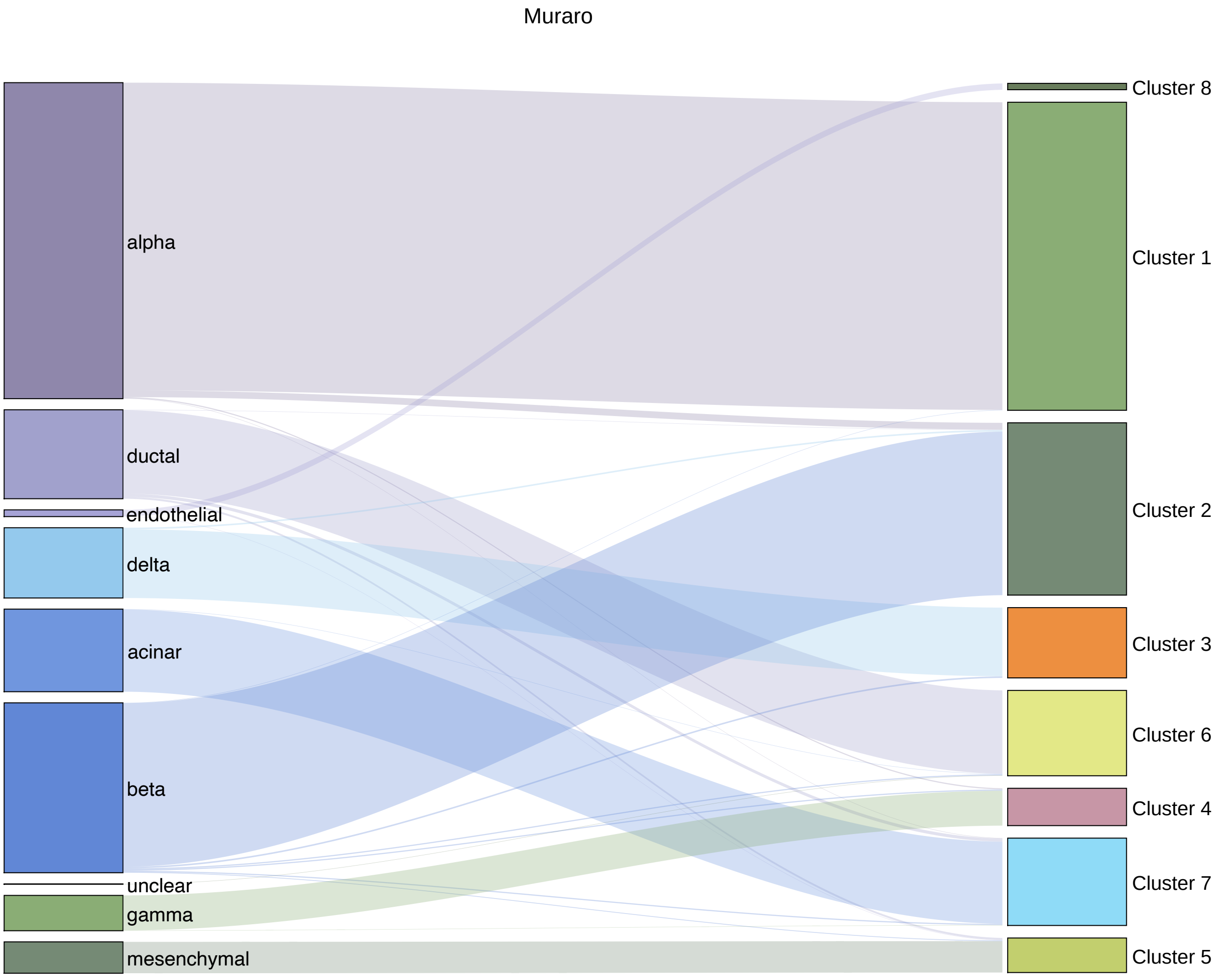

Figure S14: Sankey diagram comparison of AIGS clustering results with golden labels on Muraro [12] datasets. AIGS placed the unclear cell type into different categories, while accurately classifying other cell types, including types named alpha, acinar, beta, delta, ductal, endothelial, gamma, and mesenchymal.

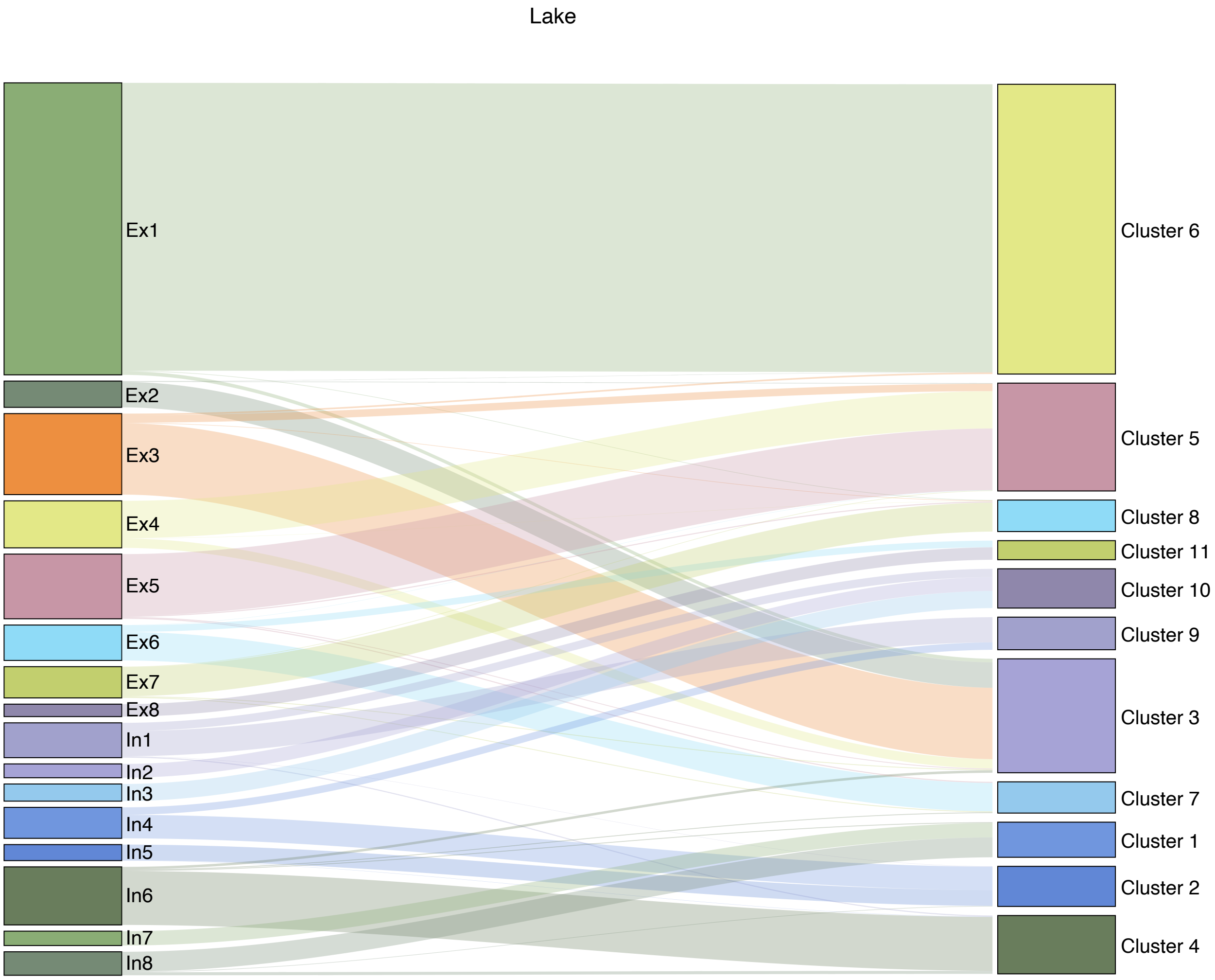

Figure S15: Sankey diagram comparison of AIGS clustering results with golden labels on Lake [13] dataset. AIGS merged Ex2 and Ex3, Ex4 and Ex5, Ex8, ln2 and ln3, ln4 and ln5, ln7 and ln8 into one category, while accurately classifying the remaining cell types, including Ex1, Ex6, Ex7, ln1, and ln6.

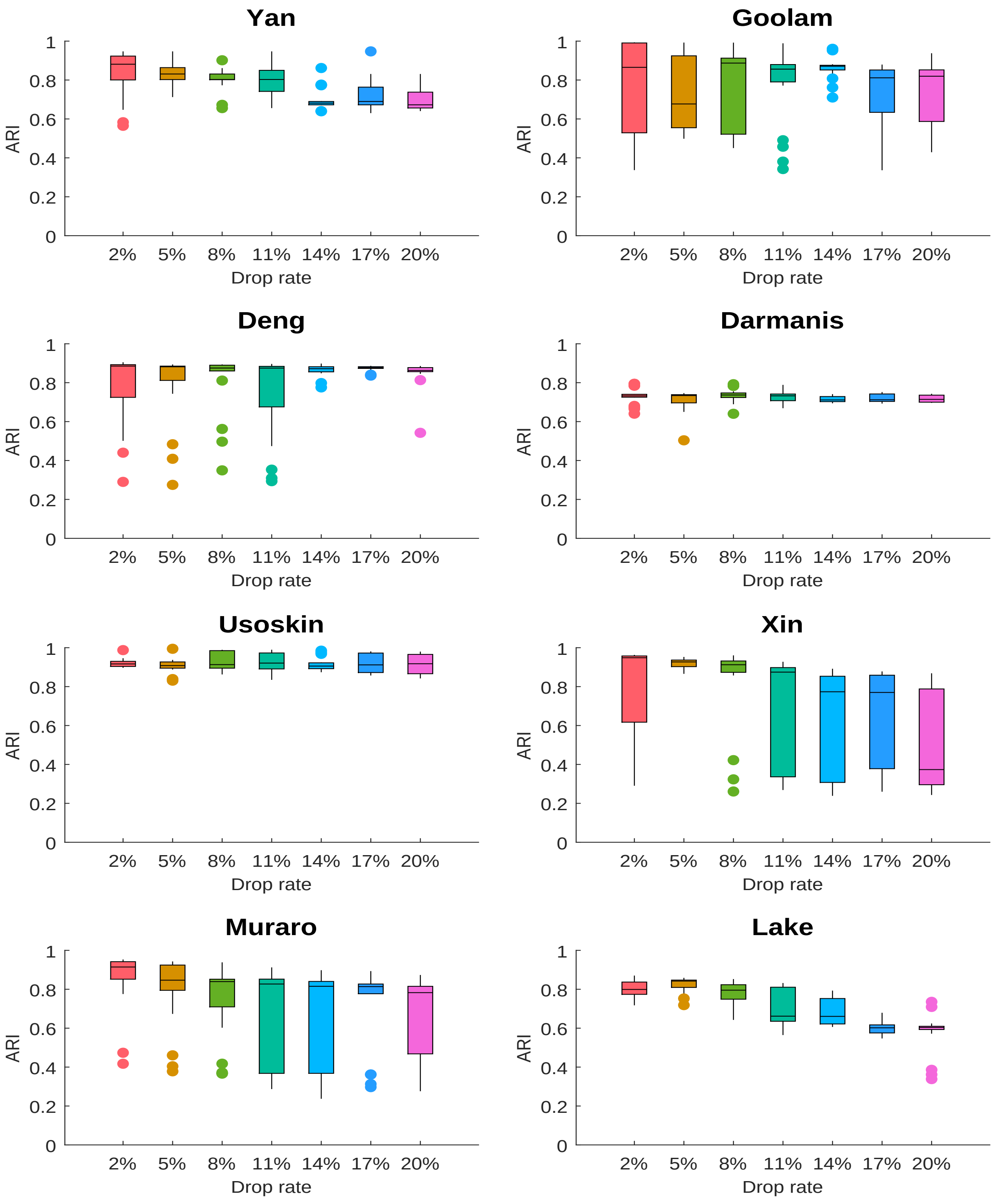

Figure S16: Box plot of Adjusted Rand Index (ARI) values for ten-time runs of AIGS with randomly altered gene expression at 2%, 5%, 8%, 11%, 14%, 17%, and 20% positions on eight datasets [6, 7, 8, 9, 10, 11, 12, 13].

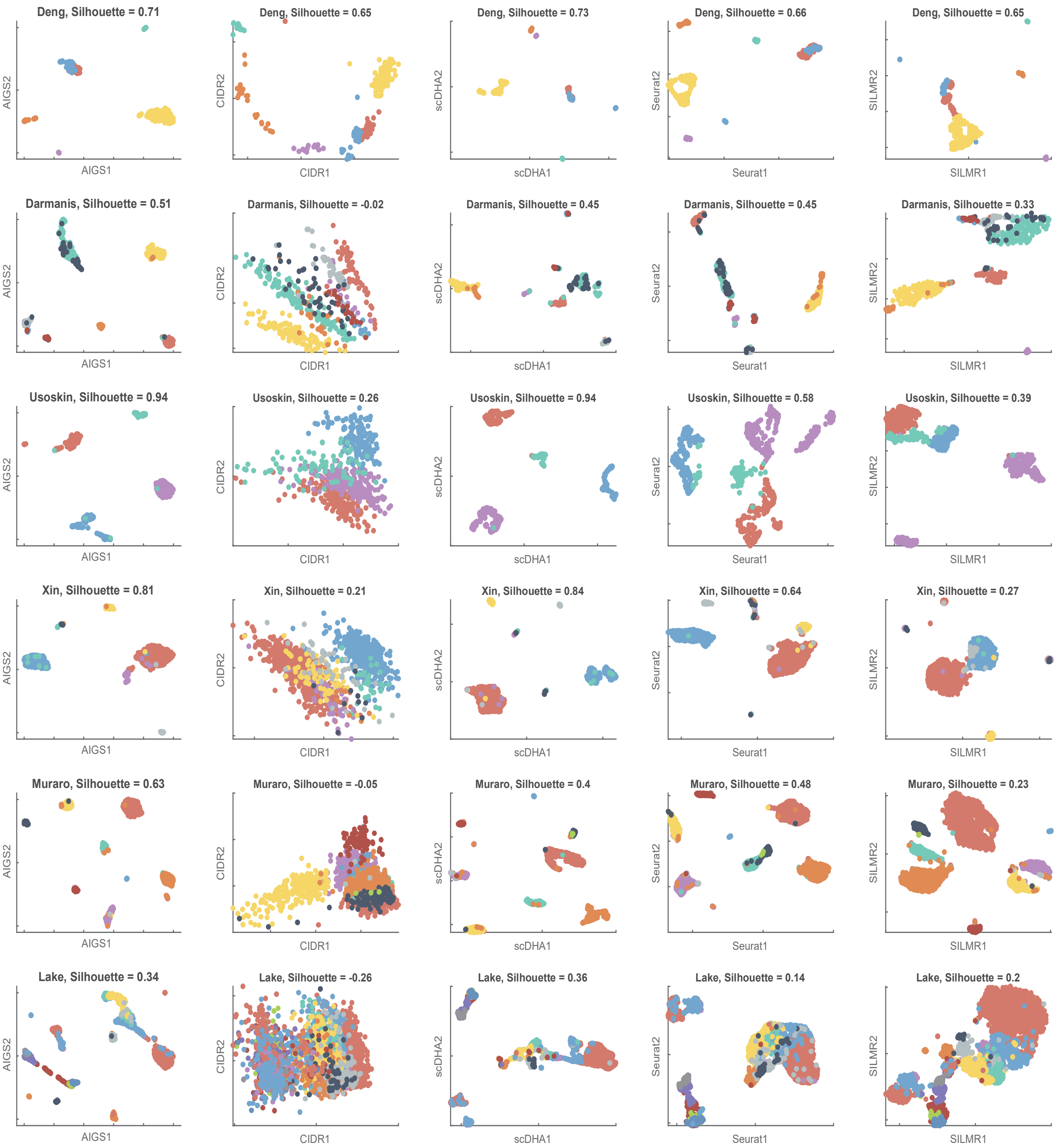

Figure S17: Visualization results comparison of AIGS, CIDR [5], scDHA [1] , Seurat [2], and SIMLR [4] (from left to right) on Deng [8], Darmanis [9], Usoskin [10], Xin [11], Muraro [12], and Lake [13] datasets (from top to bottom): AIGS's visualizations exhibit high silhouette coefficients (see **Note D.1**), indicating accurate representation of cell distribution and topological structure in high-dimensional space.

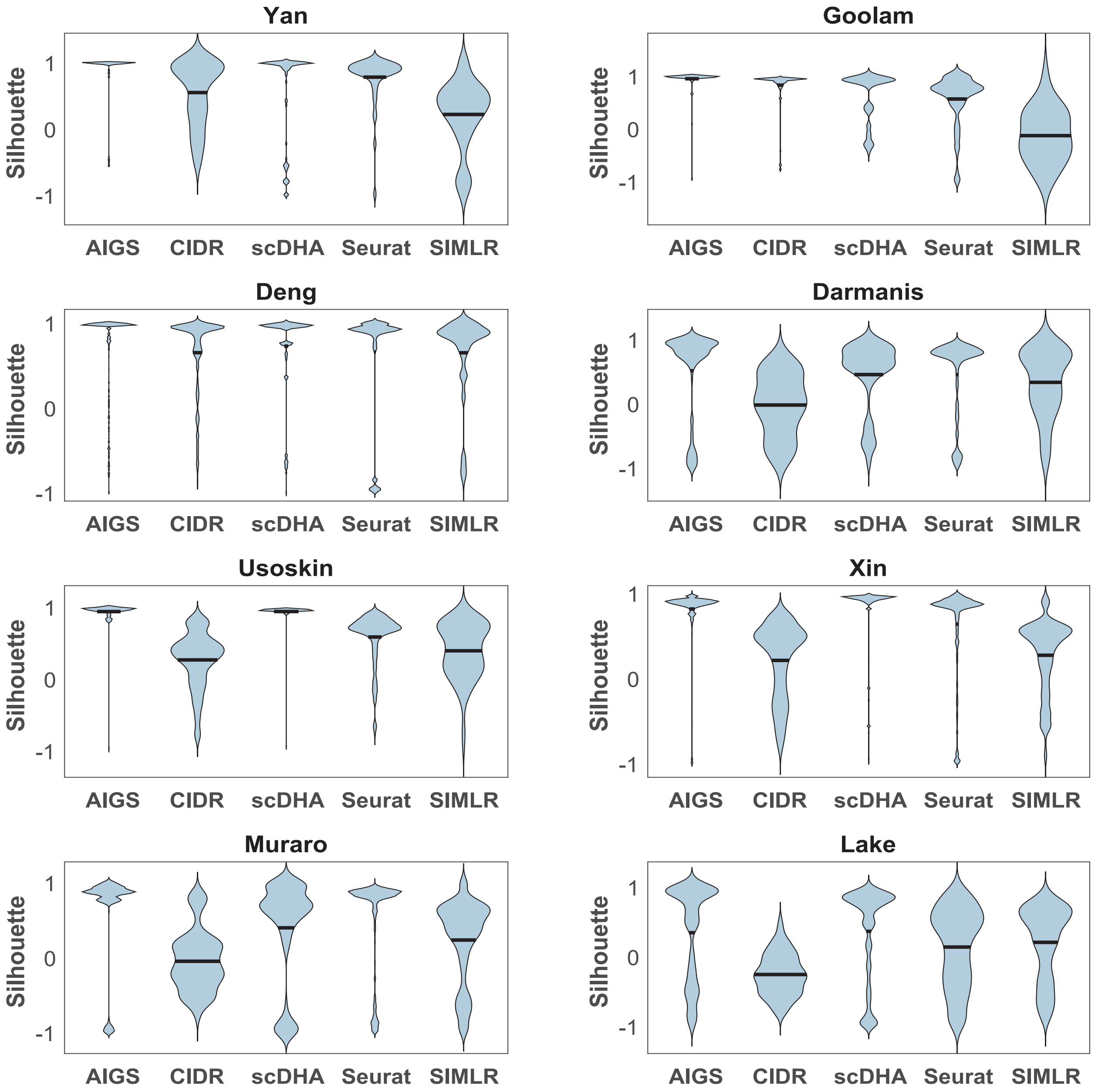

Figure S18: Violin plots of silhouette coefficients: Comparing AIGS, CIDR [5], scDHA [1], Seurat [2], and SIMLR [4] visualization results on eight datasets [6, 7, 8, 9, 10, 11, 12, 13]. The findings indicate that AIGS's visualization consistently attains more accurate and stable low-dimensional cell embeddings in comparison to the other methods.

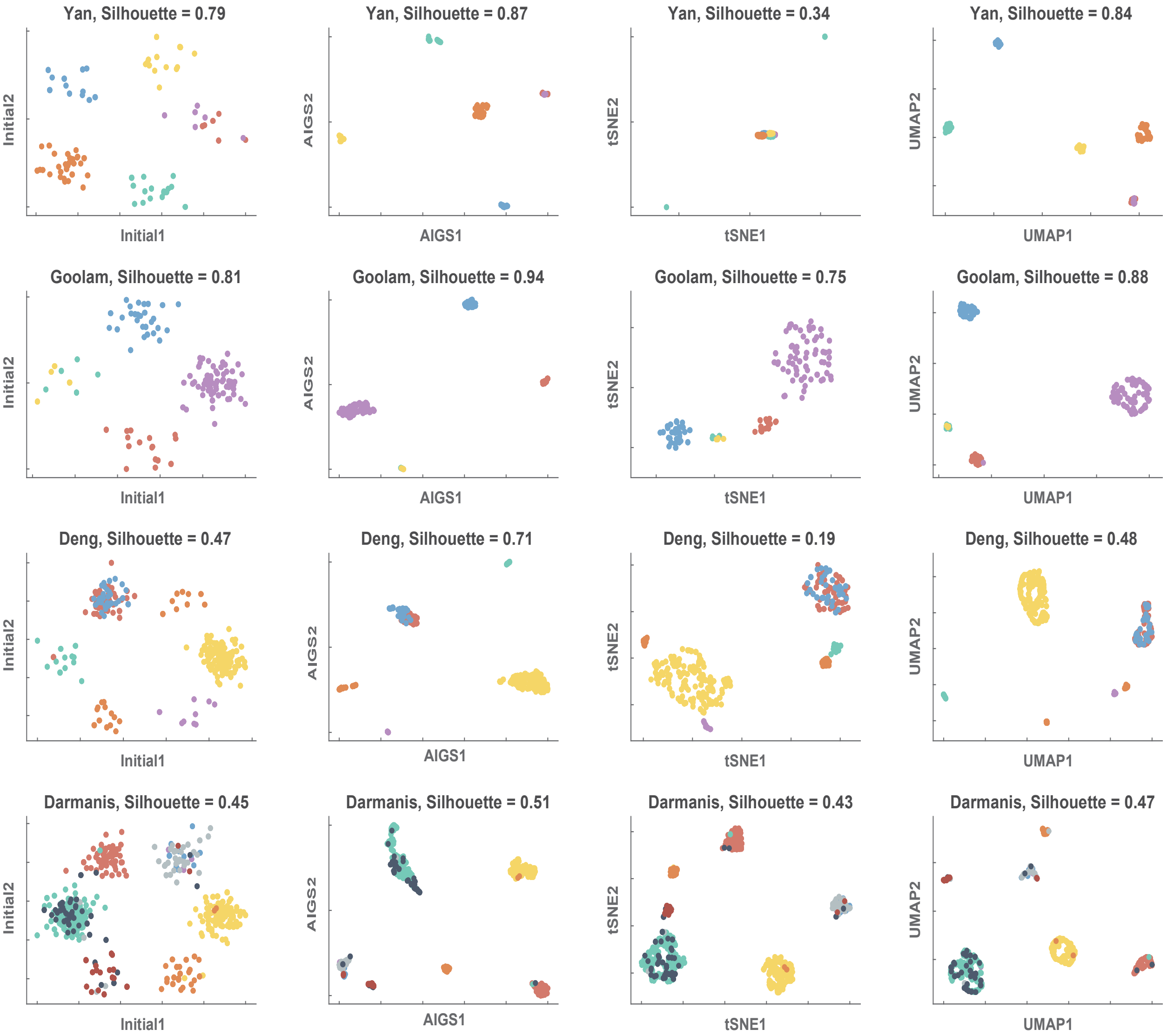

Figure S19: Comparison of silhouette coefficients for Yan[6], Goolam[7], Deng[8], and Darmanis[9] datasets using AIGS, t-SNE[28], and UMAP[30] visualizations with the same initial values. Although the initial visualizations may appear satisfactory in reflecting the features of the datasets' distribution in high dimensions, t-SNE generally reduces both intra-cluster and inter-cluster distances simultaneously, resulting in inaccurate visualizations. In contrast, AIGS and UMAP not only compress intra-cluster distances but also expand inter-cluster distances, providing more accurate reflections of the data distribution.

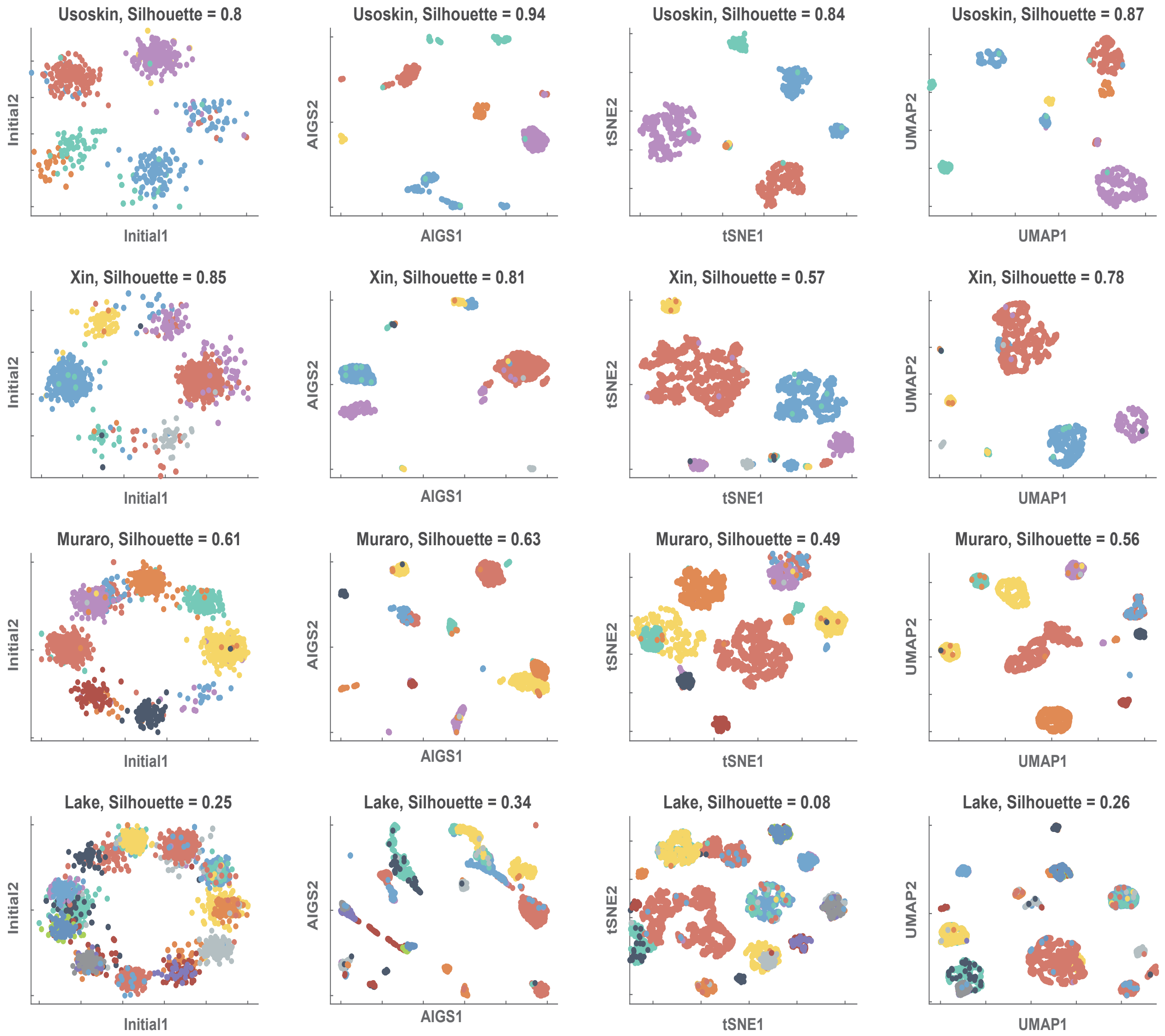

Figure S20: Comparison of silhouette coefficients for Usoskin[10], Xin[11], Muraro[12], and Lake[13] datasets using AIGS, tSNE[28], and UMAP[30] visualizations with the same initial values.

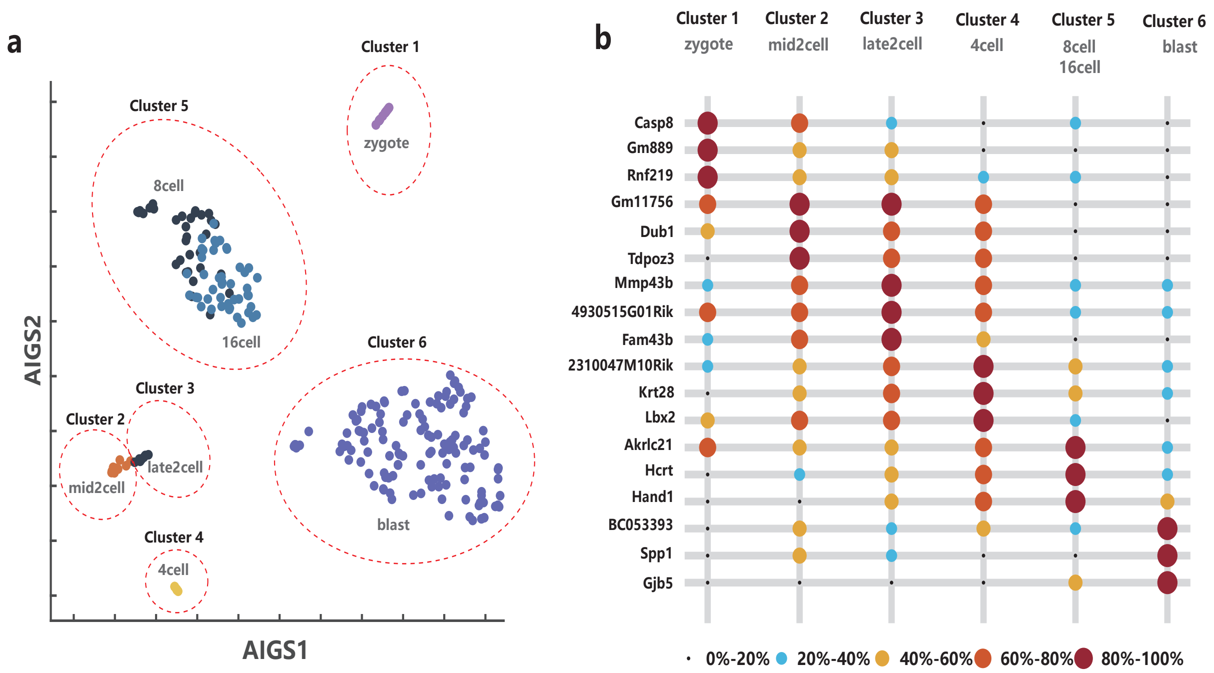

Figure S21: (a):Visualization Results of Deng Data set. Each color represents the golden labels of the Deng dataset. The red dashed line denotes the clustering results of AIGS. AIGS successfully identifies two sub-types in the 2-cell stage, namely mid2cell and late2cell. While merging the cells in the 8cell and 16cell stages into a single cluster. In addition, AIGS demonstrates accurate clustering performance in the other categories. (b): Marker gene expression across different cell types in the Deng dataset [8], top 3 markers identified for each cell type. Point size represents average expression level and colors indicate expression levels as shown in the legend. AIGS clusters match with ground truth labels and identify corresponding marker genes. The Consistent strong expression of marker genes in the 8cell and 16cell stages indicates a high degree of similarity between these two stages, highlighting their extreme resemblance. On the other hand, although the mid2cell and late2cell stages have different marker genes, their expression levels are similar. This similarity in gene expression suggests that these two stages represent closely related sub-types within the dataset.

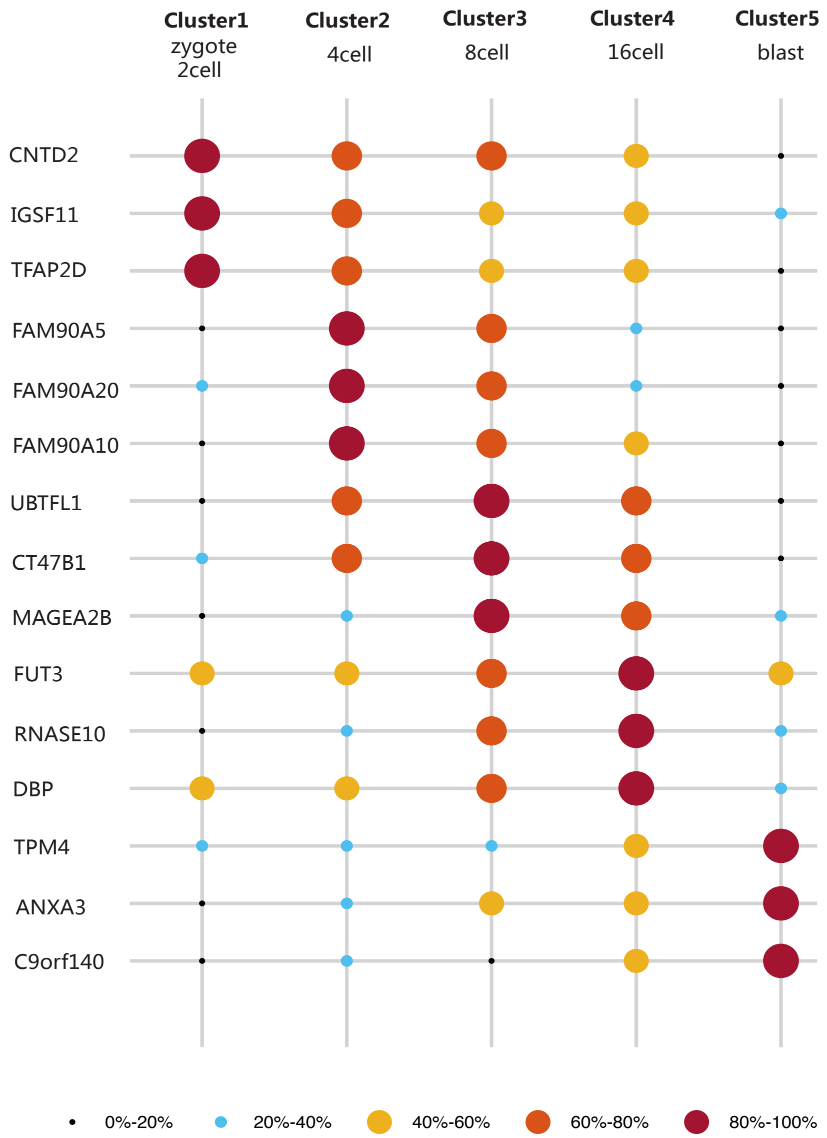

Figure S22: Marker gene expression across different cell types in the Yan dataset, top 3 markers identified for each cell type. Point size represents average expression level and colors indicate expression levels as shown in the legend. AIGS clusters match with ground truth labels and identify corresponding marker genes. AIGS clusters zygote and 2-cell stage cells into a single category (Cluster 1). Interestingly, the marker genes identified in Cluster 1 exhibit significantly different expression levels compared to the other categories. This discrepancy in gene expression suggests that Cluster 1 represents a distinct subset of cells with unique characteristics during the early developmental stages.

#### B Supplementary Table

Table 1: Description of the eight single-cell datasets used to evaluate the performance of computational methods.

| dataset | Tissue | Number of Samples | Number of Genes | Number of Classes | Reference |
|---|---|---|---|---|---|
| Yan | Human embryo | 90 | 20214 | 6 | Yan et al., 2013 [6] |
| Goolam | Mouse embryo | 124 | 41428 | 5 | Goolam et al., 2016 [7] |
| Deng | Mouse embryo | 268 | 22431 | 6 | Deng et al., 2014 [8] |
| Darmanis | Human brain | 466 | 22088 | 9 | Darmanis et al., 2016 [9] |
| Usoskin | Mouse brain | 622 | 25334 | 4 | Usoskin et al., 2015 [10] |
| Xin | Human pancreas | 1600 | 39851 | 8 | Xin et al., 2016 [11] |
| Muraro | Human pancreas | 2126 | 19127 | 10 | Muraro et al., 2016 [12] |
| Lake | Human brain | 3042 | 25051 | 16 | Lake et al.,2016 [13] |

Table 2: Evaluation of clustering performance by ACC, NMI, ARI, Fmeasure, and Jaccard (see **Supplementary Note 4**) on eight single-cell datasets [6, 7, 8, 9, 10, 11, 12, 13].

| Metric | dataset | AIGS | scDHA [1] | SEURAT [2] | DropClust [3] | SIMLR [4] | CIDR [5] |
|---|---|---|---|---|---|---|---|
| ACC | Yan | **92.94** | 77.78 | 68.89 | 85.39 | 75.56 | 80.00 |
| | Goolam | **96.58** | 90.32 | 58.06 | 53.66 | 60.48 | 70.97 |
| | Deng | 82.68 | **89.18** | 47.76 | 61.05 | 62.31 | 65.30 |
| | Darmanis | **83.71** | 77.90 | 70.17 | 81.51 | 66.52 | 66.31 |
| | Usoskin | **92.37** | 81.35 | 63.34 | 81.96 | 76.53 | 87.94 |
| | Xin | **97.89** | 95.63 | 71.56 | 68.95 | 68.00 | 74.56 |
| | Muraro | **95.89** | 94.92 | 92.94 | 71.45 | 70.84 | 45.34 |
| | Lake | **78.92** | 54.57 | 59.07 | 60.80 | 42.31 | 50.03 |
| | Average | **90.12** | 82.71 | 66.48 | 70.60 | 65.32 | 67.56 |
| NMI | Yan | **94.02** | 83.29 | 66.28 | 80.10 | 74.06 | 80.47 |
| | Goolam | **95.58** | 77.97 | 56.32 | 56.70 | 56.75 | 66.59 |
| | Deng | 83.76 | **85.94** | 64.22 | 73.25 | 39.83 | 66.93 |
| | Darmanis | 73.54 | 69.32 | 69.38 | **76.47** | 49.61 | 61.18 |
| | Usoskin | **87.55** | 79.34 | 61.32 | 60.63 | 71.86 | 75.79 |
| | Xin | **92.38** | 90.95 | 67.29 | 72.40 | 58.15 | 51.07 |
| | Muraro | **90.48** | 87.04 | 83.51 | 69.76 | 58.84 | 40.84 |
| | Lake | **77.00** | 49.80 | 64.77 | 55.40 | 22.69 | 47.16 |
| | Average | **86.79** | 77.96 | 66.64 | 68.09 | 53.97 | 61.25 |
| ARI | Yan | **97.04** | 84.18 | 69.11 | 84.83 | 62.15 | 79.78 |
| | Goolam | **99.37** | 88.31 | 42.22 | 39.26 | 47.60 | 70.45 |
| | Deng | 88.31 | **89.26** | 41.37 | 54.95 | 33.10 | 51.47 |
| | Darmanis | **77.00** | 69.78 | 65.98 | 72.08 | 45.37 | 52.83 |
| | Usoskin | **93.31** | 84.90 | 56.43 | 67.95 | 65.82 | 82.18 |
| | Xin | 96.59 | **98.31** | 63.85 | 65.47 | 54.94 | 57.20 |
| | Muraro | **94.62** | 91.27 | 90.14 | 59.96 | 67.07 | 22.40 |
| | Lake | **86.32** | 57.66 | 50.23 | 62.10 | 27.11 | 46.82 |
| | Average | **91.57** | 82.96 | 59.92 | 63.33 | 50.40 | 57.89 |
| Fmeasure | Yan | **95.29** | 84.85 | 74.37 | 86.29 | 80.03 | 83.90 |
| | Goolam | **97.46** | 88.57 | 64.20 | 62.63 | 70.21 | 81.22 |
| | Deng | 87.18 | **88.54** | 60.48 | 69.44 | 61.91 | 73.15 |
| | Darmanis | 82.26 | 77.44 | 71.85 | **82.38** | 63.26 | 69.39 |
| | Usoskin | **95.32** | 87.92 | 73.74 | 85.74 | 79.72 | 85.60 |
| | Xin | **97.78** | 95.69 | 80.37 | 77.57 | 75.76 | 75.82 |
| | Muraro | **96.51** | 94.89 | 94.12 | 79.42 | 73.09 | 48.93 |
| | Lake | **81.01** | 55.84 | 64.52 | 62.20 | 37.77 | 54.32 |
| | Average | **91.60** | 84.22 | 72.96 | 75.71 | 67.72 | 71.54 |
| Jaccard | Yan | **95.47** | 77.40 | 62.71 | 79.14 | 52.89 | 72.79 |
| | Goolam | **99.22** | 86.37 | 45.73 | 39.16 | 44.50 | 64.92 |
| | Deng | 85.78 | **86.46** | 39.46 | 50.08 | 44.61 | 46.18 |
| | Darmanis | **69.36** | 61.18 | 56.02 | 62.27 | 41.33 | 44.44 |
| | Usoskin | **90.89** | 79.66 | 48.55 | 61.27 | 60.67 | 77.79 |
| | Xin | 96.12 | **97.99** | 59.96 | 59.43 | 52.53 | 58.30 |
| | Muraro | **92.04** | 87.28 | 85.65 | 48.80 | 61.07 | 22.82 |
| | Lake | **79.79** | 49.70 | 40.78 | 51.60 | 28.81 | 38.81 |
| | Average | **88.58** | 78.26 | 54.86 | 56.47 | 48.30 | 53.26 |

Table 3: Proportion of zero gene expression in all gene expression across eight datasets [6, 7, 8, 9, 10, 11, 12, 13].

| Datasets | Yan | Goolam | Deng | Darmanis | Usoskin | Xin | Muraro | Lake |
|---|---|---|---|---|---|---|---|---|
| Proportion(%) | 45.55 | 68.52 | 60.46 | 80.77 | 84.62 | 85.63 | 73.04 | 53.70 |

Table 4: Comparison of computing time between AIGS and scDHA on eight single-cell datasets [6, 7, 8, 9, 10, 11, 12, 13].

| Methods | Yan | Goolam | Deng | Darmanis | Usoskin | Xin | Muraro | Lake |
|---|---|---|---|---|---|---|---|---|
| AIGS | 1.8 | 1.1 | 1.0 | 2.1 | 2.6 | 5.2 | 2.9 | 17.7 |
| scDHA | 73.18 | 86.98 | 91.76 | 96.59 | 95.37 | 115.82 | 128.40 | 120.84 |
| scDHA/AIGS Runtime Ratio | 41 | 79 | 92 | 46 | 37 | 22 | 44 | 7 |

Table 5: The accuracy of the marker gene found by clustering result of AIGS compared with that found by real label of cells on eight datasets [6, 7, 8, 9, 10, 11, 12, 13].

| Number of Marker Gene | Yan | Goolamn | Deng | Darmanis | Usoskin | Xin | Muraron | Lake |
|---|---|---|---|---|---|---|---|---|
| 5 | 73.33 | 72.00 | 50.00 | 35.56 | 80.00 | 75.00 | 84.44 | 80.00 |
| 10 | 78.33 | 76.00 | 65.00 | 36.67 | 87.50 | 76.25 | 87.78 | 85.63 |
| 15 | 81.11 | 76.00 | 73.33 | 46.67 | 91.67 | 77.50 | 88.15 | 90.83 |
| 20 | 80.83 | 76.00 | 78.33 | 54.44 | 93.75 | 76.88 | 89.44 | 93.13 |
| 25 | 79.33 | 77.60 | 81.33 | 60.00 | 95.00 | 76.50 | 90.22 | 94.50 |
| 30 | 80.00 | 78.67 | 85.00 | 62.96 | 95.83 | 76.25 | 91.48 | 95.42 |

Table 6: Comparison of silhouette coefficients in similarity matrix (see **Note D.2**) between AIGS and SIMLR on eight single-cell datasets [6, 7, 8, 9, 10, 11, 12, 13].

| Methods | Yan | Goolam | Deng | Darmanis | Usoskin | Xin | Muraro | Lake |
|---|---|---|---|---|---|---|---|---|
| AIGS | 0.19 | 0.16 | 0.01 | 0.20 | 0.10 | 0.14 | 0.10 | 0.22 |
| SIMLR | 0.28 | 0.23 | 0.28 | 0.36 | 0.17 | 0.18 | 0.20 | 0.32 |

Table 7: Comparison of silhouette coefficients in connectivity matrix (see **Note D.2**) between AIGS and SIMLR on eight single-cell datasets [6, 7, 8, 9, 10, 11, 12, 13].

| Methods | Yan | Goolam | Deng | Darmanis | Usoskin | Xin | Muraro | Lake |
|---|---|---|---|---|---|---|---|---|
| AIGS | 00.10 | 0.06 | 0.13 | 0.19 | 0.01 | 0.03 | 0.04 | 0.16 |
| SIMLR | 1.71 | 2.07 | 2.24 | 4.73 | 2.57 | 1.51 | 3.43 | 5.34 |

### C Supplementary Note 1: Definition of distance mentioned

The AIGS method utilizes the Spearman's rank correlation coefficient [21] in conjunction with the proposed scale-free metric to assess dissimilarities between cells. Prior to introducing Spearman's rank correlation coefficient, it is necessary to first introduce the Pearson correlation coefficient [22].

Given two vector $x = [x_1, \cdots, x_n]$ and $y = [y_1, \cdots, y_n]$, the Person correlation coefficient $r(x,y)$ between two vectors is defined as

$$r(x,y) = \frac{\sum_{i=1}^{n}(x_i - \bar{x})(y_i - \bar{y})}{\sqrt{\sum_{i=1}^{n}(x_i - \bar{x})^2}\sqrt{\sum_{i=1}^{n}(y_i - \bar{y})^2}}, \tag{1}$$

where $\bar{x}$ and $\bar{y}$ refers to the mean of $x$ and $y$. For a given vector $x$, the rank of $x$, denoted as $R(x)$, is the vector obtained by replacing each element of $x$ with its rank when the data is sorted. The Spearman correlation coefficient $\rho(x,y)$ between vectors $x$ and $y$ is defined as the Pearson correlation coefficient between their rank variables, denoted as $R(x)$ and $R(y)$, respectively. Specifically, the Spearman correlation coefficient is calculated as:

$$\rho(x,y) = s(R(x),R(y)) = 1 - \frac{6\sum_{i=1}^{n}(R(x_i) - R(y_i))^2}{n(n^2-1)}, \tag{2}$$

while the Spearman metric is defined as

$$d^{\rho}(x,y) = 1 - \rho(x,y). \tag{3}$$

Spearman correlation coefficient or Spearman metric is a useful tool for assessing the variation between cells in cell clustering because it can capture non-linear relationships between gene expression patterns. Unlike Pearson correlation coefficient, which only captures linear relationships, Spearman correlation coefficient can capture a wide range of monotonic relationships. This is particularly useful when assessing the similarity between cells, as gene expression patterns may not always follow a strict linear relationship.

In the spectral clustering, the cosine distance is used to cluster the spectral embedding by K-means algorithm. The cosine distance always belongs to the interval $[0,2]$ and it doesn't depend on the magnitudes of the vectors. For two vectors $x, y$, the cosine distance between them is

$$d(x,y) = 1 - cos(\theta) \quad \theta = \frac{\langle x,y \rangle}{\|x\|_2\|y\|_2}, \tag{4}$$

where $\langle \cdot, \cdot \rangle$ means the inner product in Euclidean space as

$$\begin{aligned} \langle x,y \rangle &= \sum_{i=1}^{n} x_i y_i, \text{ for } x,y \in \mathbb{R}^n, \\ \langle x,y \rangle &= \sum_{i=1}^{m}\sum_{j=1}^{n} A_{ij}B_{ij}, \text{ for } A,B \in \mathbb{R}^{m\times n}, \end{aligned} \tag{5}$$

and $\|x\|_2$ is the 2-norm defined as

$$\|x\|_2 = \sqrt{\sum_{i=1}^{n} x_i^2}. \tag{6}$$

### D Supplementary Note 2: Performance Metrics

#### D.1 Silhouette Coefficient

We evaluated the tightness and separation of clusters using the silhouette coefficient [20]. The silhouette coefficient measures how well a data point fits into its assigned cluster by considering the level of inter-cluster separation and intra-cluster aggregation. Given a dataset $X = \{X_1, \dots, X_n\}$, a partition of $X$ is denoted by $\{C_1, \dots, C_k\}$, and the Euclidean distance matrix $D = (D_{ij})$ of $X$ is calculated as $D_{ij} = \|X_i - X_j\|_2$, where $\|\cdot\|_2$ is the 2-norm of the vector.

For each point $X_i$, the within-cluster distance is defined as

$$a(i) = \frac{1}{\mathrm{card}\{C_m\}} \sum_{j \in C_m} D_{ij}, \quad i \in C_m, \tag{7}$$

and the distance between different classes as:

$$b(i) = \min_{\ell \neq m} \frac{1}{\mathrm{card}\{C_\ell\}} \sum_{j \in C_\ell} D_{ij}, \quad i \in C_m. \tag{8}$$

Then the profile coefficient $s(i)$ of $X_i$ is defined as:

$$s(i) = \frac{b(i) - a(i)}{\max\{a(i), b(i)\}}, \tag{9}$$

and the silhouette coefficient $s$ of $X$ as:

$$s = \frac{1}{n} \sum_{i=1}^{n} s(i). \tag{10}$$

To evaluate the effectiveness of our AIGS method in restoring cell distribution and representing the relationships between sample points after projection, we used the silhouette coefficient. The coefficient ranges from $-1$ to 1, where higher values indicate more compact clusters within the same class and greater separation between different classes. We calculated the silhouette coefficient obtained by AIGS for the 2D projection datasets $Y$ under the true labels.

Additionally, we compared the silhouette coefficient between the Spearman metric and our novel metric on scRNA-seq datasets. This comparison allowed us to use our metric to guide gene selection based on pseudo-labeling according to the distance matrix. We retained genes that are highly expressed to achieve better performance in the clustering task.

#### D.2 Silhouette Coefficient in similarity matrix and connectivity matrix

To evaluate the quality of a similarity matrix $S$ for a given dataset $X = \{X_1, \cdots, X_n\}$, we define a partition $C = \{C_1, \cdots, C_k\}$ of $X$ into $k$ classes. The similarity between two classes $\ell$ and $m$ is then computed as the sum of similarities between all pairs of points, where one point belongs to class $\ell$ and the other belongs to class $m$. We can represent this as follows:

$$a(\ell, m) = \sum_{i \in C_\ell} \sum_{j \in C_m} S_{ij} \text{ for } \ell, \mathrm{m} \in 1, \cdots, \mathrm{k}. \tag{11}$$

We use this measure to define the silhouette coefficients of the similarity matrix (SSC) as the ratio of the sum of similarities between different classes to the sum of similarities within each class:

$$\mathrm{SSC(S)} = \frac{\sum_{\ell \neq \mathrm{m}} \mathrm{a}(\ell, \mathrm{m})}{\sum_\ell \mathrm{a}(\ell, \ell)}. \tag{12}$$

Additionally, we define the connectivity matrix $\hat{S}$ as a binary matrix obtained by replacing non-zero entries in $S$ with 1. We then define the silhouette coefficient of the connectivity matrix (CSC) as $\mathrm{SSC}(\hat{\mathrm{S}})$. When a given partition $C$ is based on the true cell types in a single-cell dataset, a smaller SSC or CSC indicates greater similarity between cells of the same type and greater dissimilarity between cells of different types, and thus a more effective similarity matrix. In the **Supplementary Table 6** and **7**, compared with SIMLR [4], AIGS provides a more sparse similarity matrix, but retains a large number of similarities of the same kind that meet the real classification results.

### E Supplementary Note 3: Clustering methods

#### E.1 K-means clustering

K-means clustering [23] is a traditional and popular unsupervised machine learning algorithm used for partitioning data points into $k$ number of clusters $S = \{S_1, \cdots, S_k\}$ based on their distance, where $k$ is a predetermined number. The K-means algorithm minimizes the distance between the points in each cluster and the centroid of the cluster. Formally, the optimal solution is found as:

$$\underset{S}{\operatorname{argmin}} \sum_{i=1}^{k} \sum_{x_j \in S_i} d(x_j, \mu_i)^2, \tag{13}$$

where $\mu_i$ is the centroid of data points in $S_i$ and $d(x, \mu_i)$ is the distance between $x_j$ and $\mu_i$. The most common algorithm used for optimization is an iterative technique. Given an initial set of $\{m_1^{(1)}, \cdots, m_k^{(1)}\}$, the algorithm proceeds by alternating between two steps:

1. Assign each observation to the cluster with the nearest mean

$$S_i^{(t)} = \left\{ x_p : d(x_p, m_i^{(t)}) \leq d(x_p, m_j^{(t)}), \forall j, 1 \leq j \leq k \right\}. \tag{14}$$

2. Recalculate centroids for datasets assigned to each other

$$m_i^{(t+1)} = \frac{1}{\operatorname{card}\{S_i^{(t)}\}} \sum_{x_j \in S_i^{(t)}} x_j. \tag{15}$$

Although the K-means algorithm can converge when the assignments no longer change, it is not guaranteed to find the global optimum. The algorithm is sensitive to the initial choice of centroids and may get trapped in a local minimum. As a result, it is often recommended to run the algorithm multiple times with different initializations to increase the chances of finding a good solution.

#### E.2 Spectral clustering

Spectral clustering [24] is a technique that uses the eigenvectors of the similarity matrix to perform dimensionality reduction, called spectral embedding, before clustering into fewer dimensions. Given a set of data $\{x_1, \cdots, x_n\}$, the similarity is often defined by a Gaussian kernel function as

$$S_{ij} = \exp\left(-\frac{d(x_i, x_j)^2}{2\sigma^2}\right), \tag{16}$$

or self-tuning version

$$S_{ij} = \exp\left(-\frac{d(x_i, x_j)^2}{2\sigma_i \sigma_j}\right), \tag{17}$$

where $\sigma$ is a tuning parameter, $d(\cdot, \cdot)$ means the metric, $\sigma_i$ is the p-th smallest value in set $\{d(x_1, x_i), \cdots, d(x_n, x_i)\}$ and $\sigma_j$ is similar. However, when creating a similarity matrix, typically only the non-zero similarities between a sample and its neighborhood are taken into account.

$$S_{ij} = \begin{cases} \exp\left(-\frac{d^2(x_i, x_j)}{2\sigma_i \sigma_j}\right), & x_j \in N(x_i); \\ 0, & \text{Otherwise}, \end{cases} \tag{18}$$

where $N(x_i)$ means the $k$ neighbour of $x_i$. Generally speaking, there are two ways to select the neighborhood $N(x_i)$:

1. The first method is to choose all samples whose distance from $x_i$ is smaller than a given threshold which is independent of the distance between $x_i$ and other samples.;
2. The second method is to select the top $k$ samples that are closest to $x_i$ in terms of distance.

When the distribution of points for different class is uneven, or the density in the neighborhood of data point is different, the self tuning version definition can often better reflect the similarity between samples since the distance is scaled by that between points instead of using a fixed constant.

Then, defined the degree matrix

$$D = \operatorname{diag}\left(\sum_j s_{1j}, \cdots, \sum_j s_{Nj}\right), \tag{19}$$

and minimize the loss function

$$\min_{H^T SH=I,\ H\geq 0} \operatorname{trace}(H^T SH), \tag{20}$$

which is the NP-hard problem so that relax the Non-negative properties of matrix elements and substitute $Q = D^{1/2}H$, which can obtain the relaxed problem

$$\max_{Q^T Q=I,\ Q\in\mathbb{R}^{n\times k}} \operatorname{trace}\left(Q^T D^{-1/2} S D^{-1/2} Q\right). \tag{21}$$

This is the standard trace maximization problem [25], which can be solved by finding the matrix $Q$ that contains the last $k$ eigenvectors of $D^{-1/2}SD^{-1/2}$ as columns. For the problem of computing eigenvalues and eigenvectors of sparse matrices, the Krylov subspace iterative methods [26] are often used to accelerate the process. Let $y_i \in \mathbb{R}^k$ be the vector corresponding to the $i$-th row of $Q$. Then, the points $y_i$ are clustered using the K-means algorithm based on cosine distance.

### F Supplementary Note 4: Evaluation metrics

We used five commonly used clustering accuracy metrics, including Accuracy (ACC) [15], Fmeasure [16], Jaccard coefficient (Jaccard) [17], Adjusted Rand Index (ARI) [18], and Normalized Mutual Information (NMI) [19]. These metrics assess the similarity between the clustering results and the real label of each single-cell RNA sequencing dataset.

Given a set $S$ of $n$ elements, and two clusters or partitions of these elements, namely $X = \{X_1, \cdots, X_r\}$ and $Y = \{Y_1, \cdots, Y_s\}$ with $r \geq s$. We defined $n_{ij} = \text{card}\{X_i \bigcap Y_j\}$, $n_{i\cdot} = \sum_{j=1}^{s} n_{ij} = \text{card}\{X_i\}$, and $n_{\cdot j} = \sum_{i=1}^{r} n_{ij} = \text{card}\{Y_j\}$, where $\text{card}\{\cdot\}$ denotes the size of a set. The ACC measures the proportion of correctly assigned cells and is calculated as the average of the maximum number of cells assigned to the same cluster:

$$\text{ACC} = \frac{1}{n} \sum_{i=1}^{r} \max_{j}(n_{ij}). \tag{22}$$

The definition of Fmeasure is similar to a normalized version of Accuracy and is calculated as:

$$F = \frac{2}{n} \sum_{i=1}^{r} \max_{j} \left( n_{\cdot j} \left\{ \frac{n_{ij}}{n_{i\cdot} + n_{\cdot j}} \right\} \right). \tag{23}$$

The calculation of both the Adjusted Rand Index (ARI) and the Jaccard coefficient involves the use of the combination number $\binom{N}{2}$, which represents the total number of possible pairs that can be formed from a set of $N$ cells. Specifically, the Jaccard coefficient can be calculated as follows:

$$J = \frac{\sum_{i=1}^{r} \sum_{j=1}^{s} \binom{n_{ij}}{2}}{\sum_{i=1}^{r} \binom{n_{i\cdot}}{2} + \sum_{j=1}^{s} \binom{n_{\cdot j}}{2} - \sum_{i=1}^{r} \sum_{j=1}^{s} \binom{n_{ij}}{2}}, \tag{24}$$

and ARI is calculated as:

$$\text{ARI} = \frac{\sum_{i=1}^{r} \sum_{j=1}^{s} \binom{n_{ij}}{2} - \left[ \sum_{i=1}^{r} \binom{n_{i\cdot}}{2} \sum_{j=1}^{s} \binom{n_{\cdot j}}{2} \right] / \binom{n}{2}}{\frac{1}{2} \left[ \sum_{i=1}^{r} \binom{n_{i\cdot}}{2} + \sum_{j=1}^{s} \binom{n_{\cdot j}}{2} \right] - \left[ \sum_{i=1}^{r} \binom{n_{i\cdot}}{2} \sum_{j=1}^{s} \binom{n_{\cdot j}}{2} \right] / \binom{n}{2}}. \tag{25}$$

Finally, the definition of NMI is the normalization of Mutual Information(MI). NMI takes into account the entropy between partitions to improve the accuracy of the comparison as

$$H(X) = -\frac{1}{n} \sum_{i=1}^{r} n_{i\cdot} \log\left(\frac{n_{i\cdot}}{n}\right) \quad H(Y) = -\frac{1}{n} \sum_{j=1}^{s} n_{\cdot j} \log\left(\frac{n_{\cdot j}}{n}\right), \tag{26}$$

and the definition of NMI is

$$\text{NMI} = \frac{2 \sum_{i=1}^{r} \sum_{j=1}^{s} n_{ij} \left[ \log(n n_{ij}) - \log(n_{i\cdot} n_{\cdot j}) \right]}{n(H(X) + H(Y))}. \tag{27}$$

A higher value of the five metrics indicates a greater similarity between the two partitions. Our evaluation, which involved measuring these metrics between the clustering results generated by AIGS and the true labels of each scRNA-seq data, highlights the superiority of our technique for clustering tasks.

### G Supplementary Note 5: Description of visualization algorithm

#### G.1 Explanation on loss function and optimization process in visualization

We use the cross-entropy of Bernoulli distributions $P$ and $Q$ to measure the similarity between cells:

$$H(P,Q) = -p\log(q) - (1-p)\log(1-q),\ P(\omega) = p,\ Q(\omega) = q. \tag{28}$$

Specifically, we define $\omega$ as a pair of connected cells and the similarity between cells as the probability of the occurrence of $\omega$. Our goal is to ensure that the connection probability between cells is consistent in both two-dimensional and high-dimensional spaces. To achieve this, we use the following loss function:

$$f(Y) = \sum_{i,j=1}^{n} H(P_{ij}(\hat{C}), Q_{ij}(Y)). \tag{29}$$

where $P_{ij}(\hat{C})$ and $Q_{ij}(Y)$ are two Bernoulli distributions defined as

$$P_{ij}(\hat{C}) = \begin{cases} \hat{S}_{ij}, & \hat{c}_i \text{ and } \hat{c}_j \text{ are connected;} \\ 1-\hat{S}_{ij}, & \text{Otherwise,} \end{cases} \qquad Q_{ij}(Y) = \begin{cases} R_{ij}, & y_i \text{ and } y_j \text{ are connected;} \\ 1-R_{ij}, & \text{Otherwise,} \end{cases} \tag{30}$$

where $\hat{S}_{ij}$ and $R_{ij}$ represent the similarities between the original data points and their 2D embeddings, respectively.

To accelerate the optimization process in the AIGS visualization, the objective function is decomposed into a positive term $g(Y)$ and a negative term $h(Y)$, as shown below:

$$g(Y) = \sum_{\hat{S}_{ij}\neq 0} \hat{S}_{ij}\log\left(\frac{1}{R_{ij}}\right), \quad h(Y) = \sum_{i\neq j}(1-\hat{S}_{ij})\log\left(\frac{1}{1-R_{ij}}\right), \tag{31}$$

When working with large datasets containing a vast number of cells, it is essential to optimize the objective function efficiently, as gradient descent may be very time-consuming and may result in gradient explosion if the step size is not carefully chosen. To address this, we utilize the stochastic gradient descent algorithm [27]. Furthermore, to speed up the convergence of the random gradient descent, we employ negative sampling [28] and an alternating minimization method [29]. The gradient is computed as a combination of the positive and negative samples in each iteration of the algorithm:

$$G_g(y_i) = \sum_{j\in I_1} \nabla_{y_i}\left\{\hat{S}_{ij}\log\left(\frac{1}{R_{ij}}\right)\right\}, \quad G_h(y_i) = \sum_{j\in I_2} \nabla_{y_i}\left\{(1-\hat{S}_{ij})\log\left(\frac{1}{1-R_{ij}}\right)\right\}, \tag{32}$$

where $\nabla_{y_i}\{\cdot\}$ refers to the calculation of the gradient of the vector variable $y_i$. Additionally, the two sets denoted by $I_1$ and $I_2$ means randomly chosen positive and negative index sets:

$$\begin{aligned} I_1 &= \{j : \hat{S}_{ij} \neq 0\},\ \text{card}\{I_1\} = c_1, \\ I_2 &= \{j : i \neq j\},\ \text{card}\{I_2\} = c_2, \end{aligned} \tag{33}$$

where $c_1$ and $c_2$ are set as 20 by default, while in the visualization aimed at subtype resolution, we adjust $c_1$ and $c_2$ to 30 and 100, respectively. Then update $y_i$ to

$$y_i \leftarrow y_i - \alpha(G_g(y_i) + G_h(y_i)). \tag{34}$$

Subsequently, each $y_i$ is updated in an alternating manner during each epoch until convergence is achieved or a predetermined maximum number of epochs is reached.

#### G.2 Method for choosing parameter in visualization

In visualization, super parameters $a$ and $b$ need to be adjusted. We determine parameters by solving the optimization problem as

$$\min_{a,b} \sum_{i,j=1}^{N_t} \left\| \frac{1}{(1+a\|y_i-y_j\|_2^{2b})} - \exp\left(\|y_i-y_j\|_2^2\right) \right\|, \tag{35}$$

where $Y = [y_1, \cdots, y_{N_t}]$ is the initial value given by AIGS. However, it becomes more and more difficult to solve this optimization problem as the number of cells increases. So we turn the problem into determining parameter $a$ and $b$, so that $1/(1+ax^{2b})$ and $\exp(-x^2)$ are close on $[0, \max_Y\{\|y_i-y_j\|_2\}]$. In the program, we take the node obtained by equidistant division of interval $[0, \max_Y\{\|y_i-y_j\|_2\}]$ as $\{x_i\}$, where the number of nodes is set as 300, and solve the optimization problem to obtain parameters $a, b$.

$$\min_{a,b} \sum_{i=1}^{300} \left\| \frac{1}{1+ax_i^2} - \exp(-x_i^2) \right\|_2^2. \tag{36}$$

At this time, the algorithm achieves the large-scale extension.

#### H Supplementary Note 6: Data sources of single-cell RNA-seq

To evaluate the effectiveness of our method, we utilized eight classical datasets that collectively encompassed hundreds to thousands of distinct cell types (**Supplementary Table 1**). A comprehensive description of each single-cell RNA-seq dataset can be found below.

1. The Yan dataset [6] consists of 90 embryonic cells, including 6 distinct developmental stages, including zygotes, 2-cell, 4-cell, 8-cell, 16-cell, and blast. This dataset was specifically designed to capture the transcriptomic changes occurring throughout embryonic development. In total, the dataset generated 438 Gb of sequencing data, with an average of 35.3 million reads per cell and a read length of 100 bp.
2. The dataset generated by Goolam et al [7] includes 39 single-cell transcriptomes from various stages of embryonic development, including 2, 4, 8, 16, and 32 cells. Quality control measures were implemented to ensure reliable data, and stringent criteria were applied based on reads mapped to endogenous RNA molecules, the number of genes with at least 10 reads per million, and reads mapped to mitochondrial genes, with excluded data meeting these criteria.
3. The Deng dataset [8] consists of 269 transcripts obtained from oocytes at different developmental stages using RNA sequencing technologies. At the early stages of development, including the zygote and two-cell stage, maternal RNA comprised the majority of transcripts, with minimal contribution from the paternal genome. However, as development progressed, the proportion of maternal RNA decreased and the levels of maternal and paternal transcripts became comparable by the 4-cell stage.
4. The Darmanis dataset [9] comprises 8 anterior temporal lobe tissues obtained from adult patients during surgical resection and a set of 4 fetal cortical tissues with a gestational age of 16-18 weeks. In total, 466 cells were isolated and profiled, including major brain cells types such as astrocytes, oligodendrocytes, oligodendrocyte precursor cells, neurons, microglia, and vascular cells.
5. The Usoskin dataset [10] comprises 799 single cells from mice, including 622 neurons, 109 non-neuronal cells, and 68 cells with unknown classifications. The dataset aimed to identify previously unknown neuronal sub-types by detecting the expression of 3574 ś 2,010 genes in each cell.
6. The Xin dataset [11] comprises transcriptomes from 3709 cells isolated from 12 non-diabetic patients and 6 patients with type 2 diabetes (T2D). Following quality control measures, 1492 transcriptomes from the alpha, beta, delta, and PP cells were selected for further analysis, with a total of 29,887 genes detected.
7. The Muraro dataset [12] comprises pancreatic tissue isolated from four organ donors, which was then cultured in vitro for 3 to 5 days. Fluorescence-activated cell sorting (FACS) and single-cell sequencing were used to generate an average of 4,262 transcripts per cell, with a median of 1,958 genes detected.
8. The Lake dataset [13] comprises 3227 mononuclear neurons, including 972 inhibitory neurons and 2253 excitatory neurons, isolated from 6 cortical regions of the postmortem female brain.
